# Two Classes of DNA Gyrase Inhibitors Elicit Distinct Evolutionary Trajectories Toward Resistance in Gram-Negative Pathogens

**DOI:** 10.1101/2023.06.26.546596

**Authors:** Semen A. Leyn, James E. Kent, Jaime E. Zlamal, Marinela L. Elane, Maarten Vercruysse, Andrei L. Osterman

**Affiliations:** Sanford Burnham Prebys Medical Discovery Institute, La Jolla, California, USA; Roche Pharma Research and Early Development, Immunology, Inflammation, and Infectious Diseases, Basel, Switzerland

## Abstract

Comprehensive knowledge of mechanisms driving the acquisition of antimicrobial resistance is essential for the development of new drugs with minimized resistibility. To gain this knowledge, we combine experimental evolution in a continuous culturing device, the morbidostat, with whole genome sequencing of evolving cultures followed by characterization of drug-resistant isolates. Here, this approach was used to assess evolutionary dynamics of resistance acquisition against DNA gyrase/topoisomerase TriBE inhibitor GP6 in *Escherichia coli* and *Acinetobacter baumannii*. The evolution of GP6 resistance in both species was driven by a combination of two classes of mutational events: (i) amino acid substitutions near the ATP-binding site of the GyrB subunit of the DNA gyrase target; and (ii) various mutations and genomic rearrangements leading to upregulation of efflux pumps, species-specific (AcrAB/TolC in *E. coli* and AdeIJK in *A. baumannii*) and shared by both species (MdtK). A comparison with the experimental evolution of resistance to ciprofloxacin (CIP), previously performed using the same workflow and strains, revealed fundamental differences between these two distinct classes of compounds. Most notable were non-overlapping spectra of target mutations and distinct evolutionary trajectories that, in the case of GP6, were dominated by upregulation of efflux machinery prior to (or even in lieu) of target modification. Most of efflux-driven GP6-resistant isolates of both species displayed a robust cross-resistance to CIP, while CIP-resistant clones showed no appreciable increase in GP6-resistance.

**Importance:** The significance of this work is in assessing the mutational landscape and evolutionary dynamics of resistance acquisition against a novel antibiotic, GP6. This approach showed that, in contrast to ciprofloxacin (CIP), a previously studied a canonical DNA gyrase/topoisomerase-targeting clinical antibiotic, evolution of GP6-resistance is driven largely by early and most prominent mutational events leading to upregulation of efflux machinery. An identified asymmetry in cross-resistance of evolved GP6- vs CIP-resistant clones provides important guidelines for rational selection of potential treatment regimens. This study illustrates the utility of the established morbidostat-based comparative resistomics workflow for the assessment of new drug candidates and clinical antibiotics.

## Introduction

DNA gyrase GyrA/B and topoisomerase IV ParE/C are bacterial type II topoisomerases. Both enzyme complexes are vital for DNA replication, transcription and decatenation of daughter chromosomes in bacteria and have significant structural and sequence differences from the human type II topoisomerases ^1^. These proteins are established targets for common clinical antibiotics of the fluoroquinolone class ^2^ and for new drug discovery campaigns ^3,4^. The mechanism of action of fluoroquinolones involves binding at the topoisomerase-DNA interface, which allows DNA cleavage while suppressing the ligation reaction, thus introducing multiple deleterious nicks in the chromosome ^5^. Despite excellent efficacy against a broad spectrum of bacterial pathogens, the clinical utility of fluoroquinolones is hampered by rapidly spreading resistance driven largely by amino acid substitutions in a drug-binding site of its cognate targets that do not impair their enzymatic functions ^6^. These limitations of fluoroquinolones prompted numerous efforts to develop new type II topoisomerase inhibitors with a different binding mode and mechanism of action to avoid cross-resistance ^7^. Thus, Novobiocin, an aminocoumarin class antibiotic targeting the ATP-binding site of GyrB, was introduced into clinical practice in the 1960s, but discontinued in 1999 due to insufficient efficacy and safety ^8^. Two non-quinolone antibiotics have reached Phase III of clinical trials: (i) Zoliflodacin, a first-in-class spiropyrimidinetrione antibiotic ^9^; and (ii) Gepotidacin, which belongs to a novel class of triazaacenaphthylenes ^10^. These drugs bind in different sites at topoisomerase-DNA interface showing no cross-resistance with fluoroquinolones. A novel tricyclic class of pyrimidoindole GyrB/ParE inhibitors (TriBE inhibitors) active against a broad range of Gram-negative bacterial pathogens was introduced by Trius Therapeutics in 2013 ^11^.

The goal of this study was to assess the mutational landscape and evolutionary dynamics of resistance acquisition against the TriBE inhibitor GP6 ^12^ in the model system of *Escherichia coli* and in the important nosocomial pathogen *Acinetobacter baumannii*. This knowledge is essential for fundamental mechanistic understanding and for a comparative assessment of bacterial resistibility to this novel drug candidate. To this end, we applied a workflow combining experimental evolution in a continuous culturing device, the morbidostat, with high-coverage whole genome sequencing (WGS) of evolving bacterial populations followed by genotype-to-phenotype characterization of selected drug-resistant clones. This workflow (introduced in a model study of triclosan ^13^ and schematically illustrated in **Supplementary Figure S1**) was recently applied to the experimental evolution of resistance of three Gram-negative bacterial species to the most common fluoroquinolone drug ciprofloxacin (CIP) ^14^. This study revealed shared evolutionary trajectories toward CIP resistance driven by a limited set of missense mutations in the GyrA/B target (Stage I) followed by additional mutations leading to upregulation of species-specific efflux pumps (Stage II). In *A. baumannii*, (but not in *E. coli*), additional Stage II mutational events included amino acid substitutions in the secondary target ParC/E that also led to a substantial increase of the MIC of CIP (MIC^CIP^). This work provided a technological and conceptual foundation for a comparative resistomics assessment of a non-quinolone GyrB/ParE inhibitor GP6 reported here.

The performed analysis revealed fundamental mechanistic differences between these two distinct classes of type II topoisomerase inhibitors. Observed differences encompass non-overlapping spectra of target mutations and distinct evolutionary trajectories, dominated by deregulation of the efflux machinery in Stage I of evolution of resistance to GP6, prior to or even in lieu of target mutations. These differences also manifest in asymmetric cross-resistance profiles. While selected GP6 resistant (GP6^R^) clones displayed a comparably increased resistance to CIP, none of the previously characterized CIP resistant (CIP^R^) clones showed any increase in the MIC of GP6 MIC^GP6^. Overall, this study illustrates the utility of the morbidostat-based comparative resistomics approach for the assessment of resistibility of new drug candidates.

## Methods

### Bacterial strains and media

A sterile-filtered (0.22 µm) cation-adjusted Mueller-Hinton broth (MHB) with 2% dimethyl sulfoxide (DMSO) was used as a base media for both experimental evolution in morbidostat and MIC measurements. For morbidostat runs, the autoclaved antifoam SE-15 (Sigma) diluted 1/50 with water was added (to a final dilution 1/2,500). Bacterial strains used in this study were *Escherichia coli* BW25113 and *Acinetobacter baumannii* ATCC 17978.

### Test drugs

The 10 mM stock solution of GP6 compound (4-[(1R,4R,5R)-5-amino-2-azabicyclo[2.2.1]heptan-2-yl]-6-fluoro-N-methyl-2-(2-methylpyrimidin-5-yl)oxy-9H-pyrimido[4,5-b]indol-8-amine) ^12^ in absolute DMSO for all studies was diluted to a desired concentration in MHB base media maintaining 2% final DMSO concentration. GP6 was synthesized in-house as described in WO2014043272A1 (average purity 97%). Ciprofloxacin (Sigma) for orthogonal MIC measurements of GP6 resistant clones was prepared as a 10 mg/mL solution in water and used in serial dilutions with MHB/DMSO base media.

### Experimental evolution in the morbidostat

For the experimental evolution under GP6 stress we used the morbidostat device, custom-engineered based on general principles introduced by Toprak *et.al* ^15,16^. Morbidostat-based experimental evolution of resistance to GP6 for both *E. coli* and *A. baumannii* was performed using the general protocol as previously described for CIP ^14^. Both the morbidostat setup and programming are briefly described in **Supplementary Methods**.

Two morbidostat runs were performed (one for each strain) upon inoculation of all 6 glass reactors with 6 individually prepared log-phase cultures derived from glycerol stocks of 6 individual colonies of each strain (same as in the CIP study ^14^) at starting OD_600_=0.02 (20 mL in MHB/DMSO). During the run, the GP6 concentration in the drug-containing feed bottle was increased in three-five steps following the dynamics of evolving drug resistance. For *E.coli* these steps were: (i) 3.125 µM (or 2.5x MIC as compared to MIC =1.25 µM for the unevolved parental strain during 0-79 hrs of the run corresponding to ∼0.5xMIC drug concentration increment at each addition); (ii) 12.5 µM (10xMIC for 79-99 hrs); and (iii) 25 µM (20xMIC for 99-189 hrs). For *A. baumannii*: (i) 0.195 µM (1.25xMIC as compared to MIC =0.156 µM for the unevolved parental strain during 0-24hrs; ∼0.25xMIC drug concentration increment at each addition); (ii) 0.78 µM (5xMIC, for 24-51 hrs); (iii) 1.56 µM (10xMIC for 51-77 hrs); (iv) 6.24 µM (40xMIC for 77-103 hrs); and (v) 25 µM (160xMIC for 103-166 hrs). OD/drug profiles and sample collection schedules (at least 20 hrs apart) for both runs are shown in Supplementary Figure S1. These samples (10 mL each) were used to prepare cell pellets for population genomic sequencing and glycerol stocks for further clonal analysis. To account for a theoretical possibility of mutational adaptation to morbidostat experimental conditions, we have performed continuous culturing of both species in a non-selective turbidostat mode (each in one reactor for over 40 - 50 hrs) with periodic dilutions only by drug-free media followed by deep sequencing (mean coverage 5958x for *A. baumannii* and 3721x for *E.* coli) of the final populations (see **Supplementary Table S4** for details and sequencing results).

### Whole genome sequencing (WGS)

The Illumina-based WGS was used initially for selected samples of evolving bacterial population from the morbidostat runs and at the next stage of the workflow (**Supplementary Figure S1**) for the analysis of individual clones. Genomic DNA was extracted using the GenElute bacterial genomic DNA kit (Sigma-Aldrich) by the manufaturer’s protocol NA2110 for Gram-negative bacteria. Libraries for sequencing were prepared using a NEBNext Ultra II FS DNA library prep kit for Illumina modules E7810L and E7595L (New England BioLabs) by the manufacturer’s protocol (DNA input ≥100 ng) but using TruSeq DNA PCR-free CD index 20015949 (Illumina) adapters to eliminate PCR amplification. Size selection and cleanup were performed using magnetic beads AMPure XP (Beckman Coulter). Prepared libraries were quantified using a NEBNext Library Quant kit for Illumina E7630L (New England BioLabs) and pooled with volumes adjusted to normalize concentrations aiming for ∼700-fold genomic coverage for population samples or ∼200-fold genomic coverage for clones. Pooled library size and quality were analyzed with the 2100 Bioanalyzer instrument (Agilent). Pooled DNA libraries were sequenced by Novogene Co. using HiSeq 4000 (*A. baumannii* clones) or HiSeq X10 (all other samples) instruments (Illumina) with paired-end 150-bp read length.

Nanopore long-read sequencing was used to verify genomic rearrangement events (IS inserts, loci amplification/deletion) in selected clones. DNA samples were prepared using the SQK-RBK004 rapid barcoding sequencing kit (Oxford Nanopore Technologies) and SQK-LSK109 ligation sequencing kit with native barcoding expansion kit EXP-NBD104 according to the manufacturer protocol. Sequencing was performed using a MinION Mk1B and flow cell FLO-MIN106. The detailed description of our methodology for sequencing data analysis, variant calling and ranking is provided in **Supplementary Methods** along with the methods used to support population WGS data deconvolution for evolutionary dynamics and clonal analysis.

### MIC measurements

MIC values were measured for starter and evolved clones using broth dilution method specified in EUCAST standard protocols ^17^ using the same growth media base as in morbidostat-based experiments . The fresh colonies were resuspended in MHB medium and inoculated into a series of wells in 96-well plates containing 2-fold increasing concentrations of GP6 (or CIP for selected nonredundant subsets of clones). Measurements were performed by constant growth or end-point monitoring of OD_600_ in a BioTek ELx808 plate reader.

### RT-qPCR for mdtK gene expression assessment

For RT-qPCR experiments, selected GP6^R^ clones of *E. coli* BW25113 and *A. baumannii* ATCC 17978 as well as their respective unevolved parental strains were grown overnight in CA-MHB to a density of ∼ 2 x 10^8^ CFU/mL in triplicates). RNA isolation from flash-frozen pellets followed by RT-qPCR using mdtK- and gyrB-specific primers was performed as described in **Supplementary Methods.**

### Data availability

Clonal and population sequencing data are available in the SRA database by BioProject accession number PRJNA857658 (https://www.ncbi.nlm.nih.gov/bioproject/PRJNA857658); the *A. baumannii* ATCC 17978 reference genome is available in the European Nucleotide Archive by sample accession number ERS4228590 (http://www.ebi.ac.uk/ena/data/view/ERS4228590). The reference genome for *E.coli* BW25113 (PATRIC ID: 679895.18) was downloaded from the PATRIC database ^18^.

## Results and Discussion

We used a morbidostat-based experimental evolution workflow ^13,14^ illustrated in **Supplementary Figure S1** to assess and compare mutational landscapes, dynamics, mechanisms, and extent of acquired resistance to GP6 drug in two Gram-negative species, *E. coli* BW25113 and *A. baumannii* ATCC 17978.

### Mutational landscape and dynamics of GP6 resistance evolution in morbidostat

In both performed morbidostat runs, the cultures of *E. coli* BW25113 and *A. baumannii* ATCC 17978 were the subject of continuous growth in 6 parallel bioreactors over 189 and 166 hours, respectively. During these runs, the cultures were regularly diluted with drug-free or drug-containing media, depending on the observed growth rate/adaptation to gradually increasing GP6 concentration, up to ∼25 µM (**Supplementary Figure S2**). Of the periodically collected samples of evolving bacterial populations, 24 samples from the *E. coli* run (4 time points: 53, 72, 172 and 189 hrs from 6 reactors) and 18 samples from the *A. baumannii* run (3 time points: 51, 77 and 123 hrs from 6 reactors) were a subject of Illumina WGS.

Mutational variants mapped in these samples were based on comparison to corresponding reference genomes ^14^ and included single nucleotide variants (SNVs), short indels, insertion of IS elements, and copy number variants (CNVs) manifest as deletions and amplifications of multi-gene loci. All reliably detected variants, their features and inferred consequences (such as amino acid substitutions or gene disruptions) along with their frequency (relative abundance, %) in the analyzed samples (except for CNVs expressed as averaged loci multiplicity) are captured in **Supplementary Tables S1A and B**. We ranked significant variants (potentially contributing to acquired drug resistance) by two major criteria: (i) maximal abundance across all samples (A_max_≥10%); and (ii) the number of independent occurrences per gene (N_all_≥2) combining both, identical and distinct events across all six reactors as described in **Supplementary Methods**.

The most prominently implicated genes (9 in *E. coli* and 7 in *A. baumannii)* and variants detected therein are listed in **Table 1**, alongside those implicated by our previous study on evolution of CIP resistance ^14^. The dynamics of emergence, expansion and, in some cases, shrinking (due to competition under increasing drug pressure) of these major variants are reflected in cumulative area plots in **Supplementary** Figures 3A and 3B. In both species, all major mutational events fell into two major categories: (i) drug target modification - missense mutations affecting a narrow set of amino acid residues near the respective drug-binding sites and, at least partially, shared between both species (but not between the two drugs); and (ii) efflux deregulation - various types of mutations presumably leading to upregulation of genes encoding species-specific efflux pumps, largely (but not completely) overlapping between the two drugs. A comparison of simplified cumulative area plots reflecting evolutionary dynamics of target modification vs efflux upregulation (**Figure 1**) illustrates notable differences in the order of appearance and prevalence of these two major types of mutational events during the experimental evolution of resistance to GP6 vs CIP.

**Figure 1.**
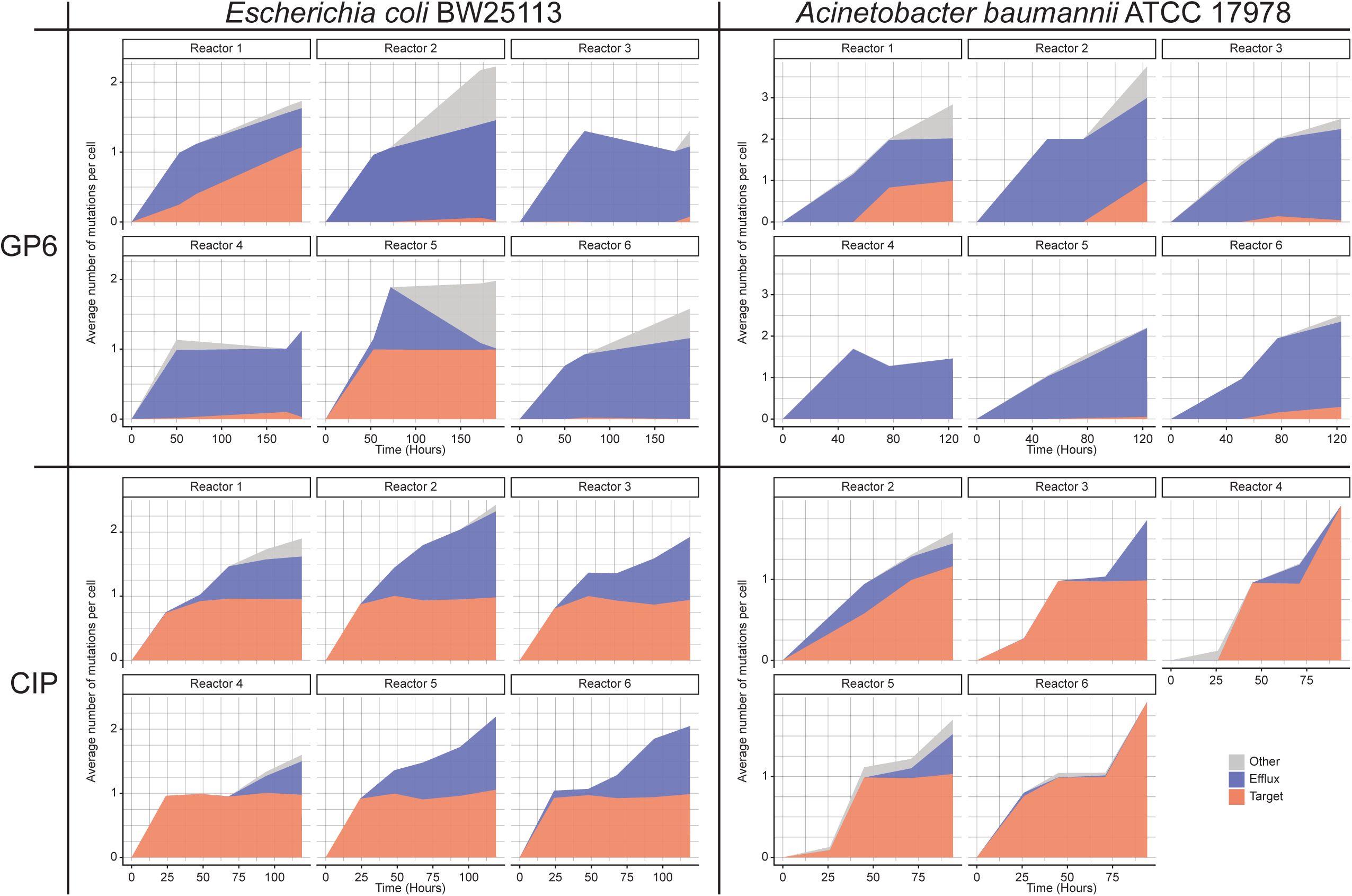
Cumulative area plots reflecting the evolutionary dynamics of the two major types of driver mutations: (i) target modifications (red area) and (ii) mutations leading to efflux upregulation (blue area); and other potentially relevant mutations (grey area). In a simplified form, the plots represent WGS data obtained for population samples in time series of collected during experimental evolution of resistance to GP6 in *E. coli* and *A. baumannii* in this study vs CIP from the previous study ^14^. The variant abundance data corresponding to each of the 6 reactors (5 in case of *A. baumannnii*_CIP) were summed up by the three categories and plotted as the average number of mutations per cell (Y-axis) vs time (X-axis). The plots reflect combined relative abundances of SNV and IS insertions, but not CNVs (loci amplifications or deletions) that cannot be accurately quantified in population WGS data (reflected in separate plots in Supplementary Figure S3).

**Table 1.** Main drivers of evolved resistance of *E. coli* and *A. baumannii* to GP6 compound and comparison with CIP resistance drivers identified in the previous study ^14^.

Of note, none of the implicated genes (as well as any other genes) acquired any appreciable mutational events in the drug-free evolutionary runs of either species in turbidostat mode (see **Supplementary Table S4**) arguing against their potential role in adaptation to non-selective continuous culturing conditions.

### Drug target modifications

As anticipated from the known mechanism of GP6 action as a GyrB inhibitor targeting the ATP-binding site, we observed amino acid substitutions (and no disruptive mutations) located in a compact area consistent with the binding mode of GP1, a structural analog of GP6 (**Figure 2**)^19^. In contrast with CIP resistance, no GyrA variants emerged on the path to GP6 resistance in either species. Likewise, no overlap was observed with previously identified CIP-resistant *E. coli* GyrB variants (**Table 1**). Moreover, despite dual-targeting characteristics of GP6 as a GyrB/ParE inhibitor, no ParE mutational variants were observed at any point in our study.

**Figure 2.**
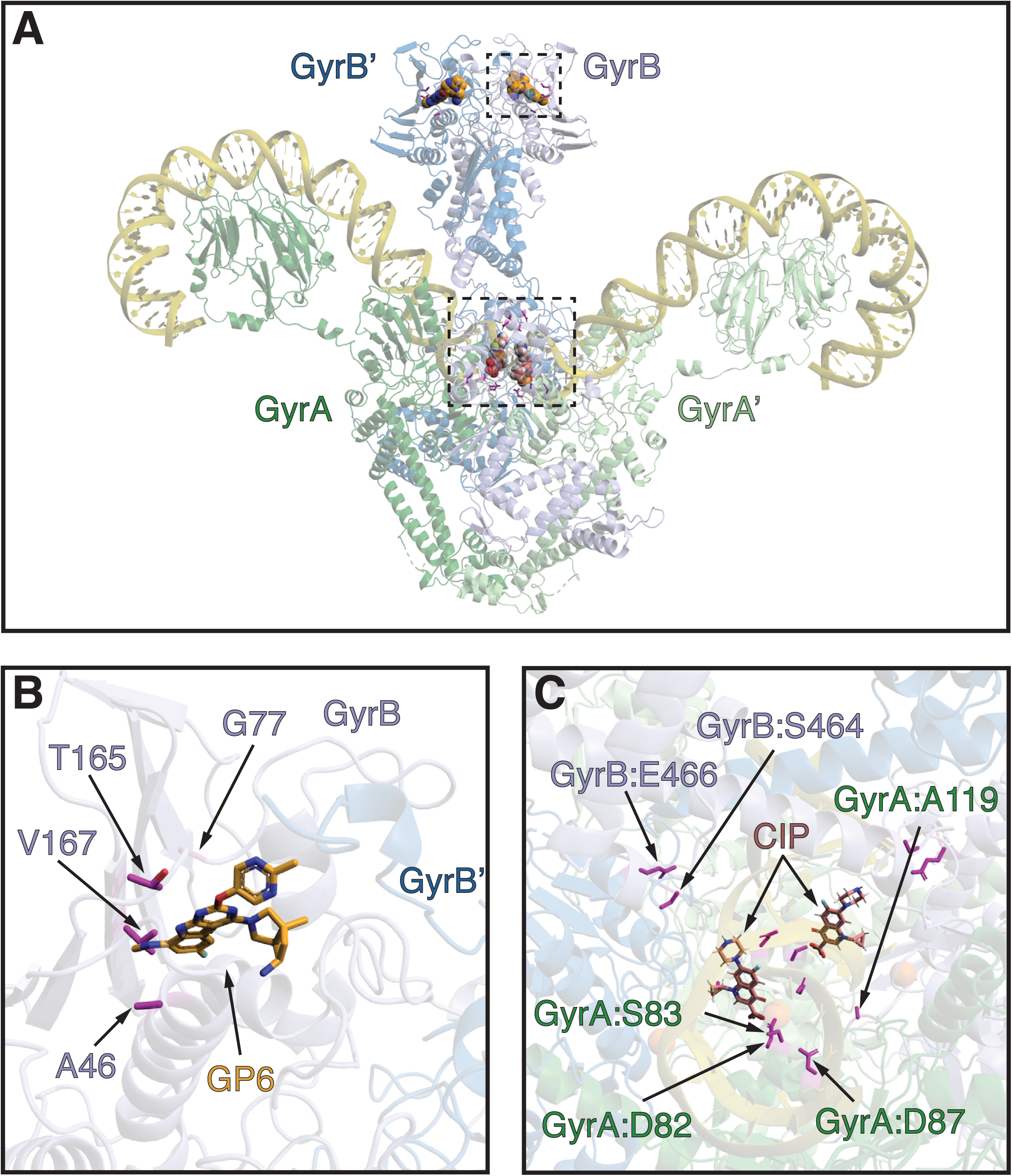
Three-dimensional projection of detected mutations in DNA gyrase complex conferring resistance to CIP or GP6 drugs. A) The structure of *E. coli* DNA Gyrase complex with DNA (PDB ID: 6RKW). GyrA subunits are colored shades of green; GyrB subunits are colored shades of blue; DNA is colored yellow. The location of GP6 (orange, modeled from 4KSG) and CIP (pink, modeled from PDB ID: 5BTC) binding sites are depicted by spheres. The amino acid residues (colored magenta) detected as mutational variants during the morbidostat-based evolution of resistance to GP6 or CIP are also shown. The areas outlined by two dashed boxes are expanded to focus on (B) GP6 and (C) CIP (variants present in the second molecule of the homodimer are not labeled).

The most remarkable differences between the two classes of DNA topoisomerase inhibitors were revealed by the comparison of evolutionary dynamics (see Figure 1 and Supplementary Figures S3AB). Indeed, in the case of CIP, GyrAB mutational variants emerged in all six reactors at the earliest stage of morbidostat-based evolution ^14^. They dominated bacterial populations throughout the entire run, combining with secondary efflux-related mutations (or, alternatively, with ParC variants as in *A. baumannii*) only at a later stage under stronger drug pressure. This is in contrast with evolutionary dynamics observed for GP6 where the early emergence of GyrB variants was observed exclusively in *E. coli* and only in 2 out of 6 bioreactors (Supplementary Figure S3A). In the case of *A. baumannii*, GyrB variants emerged only at a later stage, after substantial accumulation of efflux-upregulating mutations, reaching high abundance in 2 out of 6 bioreactors (Supplementary Figure S3B).

The observed evolutionary trajectories may be driven by different factors and combinations thereof. Thus, it is possible that the similar affinity of GP6 for GyrB and ParE may hinder the emergence of a bacterial population containing only GyrB ATP-binding site mutations, as such events would not be expected to provide a substantial advantage under GP6 pressure ^11^. (That said, the exclusive appearance of GyrB rather than ParE, variants highlights the slight difference in GP6 affinity for these two targets ^11^). Moreover, without engineering and characterizing the corresponding clean mutants, we cannot rule out the possibility that *gyrB* (or *parE*) mutations do not provide sufficient drug resistance or have some epistatic interactions with mutations in other genes. It is also conceivable that mutations in the GyrB ATP-binding site (required for GP6 resistance) in contrast with the mutations at the GyrAB-DNA interface (driving CIP resistance) have higher fitness costs than those causing efflux upregulation.

### Efflux deregulation

The overwhelming majority of the identified significant mutational events driving GP6 resistance in both species affect genes regulating their efflux machinery (**Table 1** and **Figure 3)**. In *E. coli*, the most prominent are IS insertions in the coding region of AcrR, a negative transcriptional regulator of the AcrAB efflux pump ^20^. Multiple AcrR disruptive variants are observed in all 6 bioreactors, reaching A_max_∼100% in two reactors (see **Supplementary Figure S3A**). Other *E. coli* variants associated with AcrAB regulation included: (i) missense mutations AcrR:Thr5Asn and AcrR:Thr5Ile; and (ii) distinct noncoding variants in the intergenic region of the *acrR*<>*acrAB* divergon, overlapping with the known AcrR binding site ^20^. Several intergenic mutational events were observed upstream of the *marR* gene including large deletions spanning SD sequence and, in one case, a start codon. These loss-of-function events are expected to derepress transcription of the *marRA* operon, which would in turn upregulate the expression of AcrAB and TolC components of the *E. coli* efflux machinery. Similar *acrR* and *marR* mutations were detected in our CIP study ^14^, but in contrast with it, on the path to GP6 resistance, we observed no *soxR* mutations that would increase AcrAB/TolC expression via derepression of *soxS* gene (**Table 1** and **Figure 3**).

**Figure 3.**
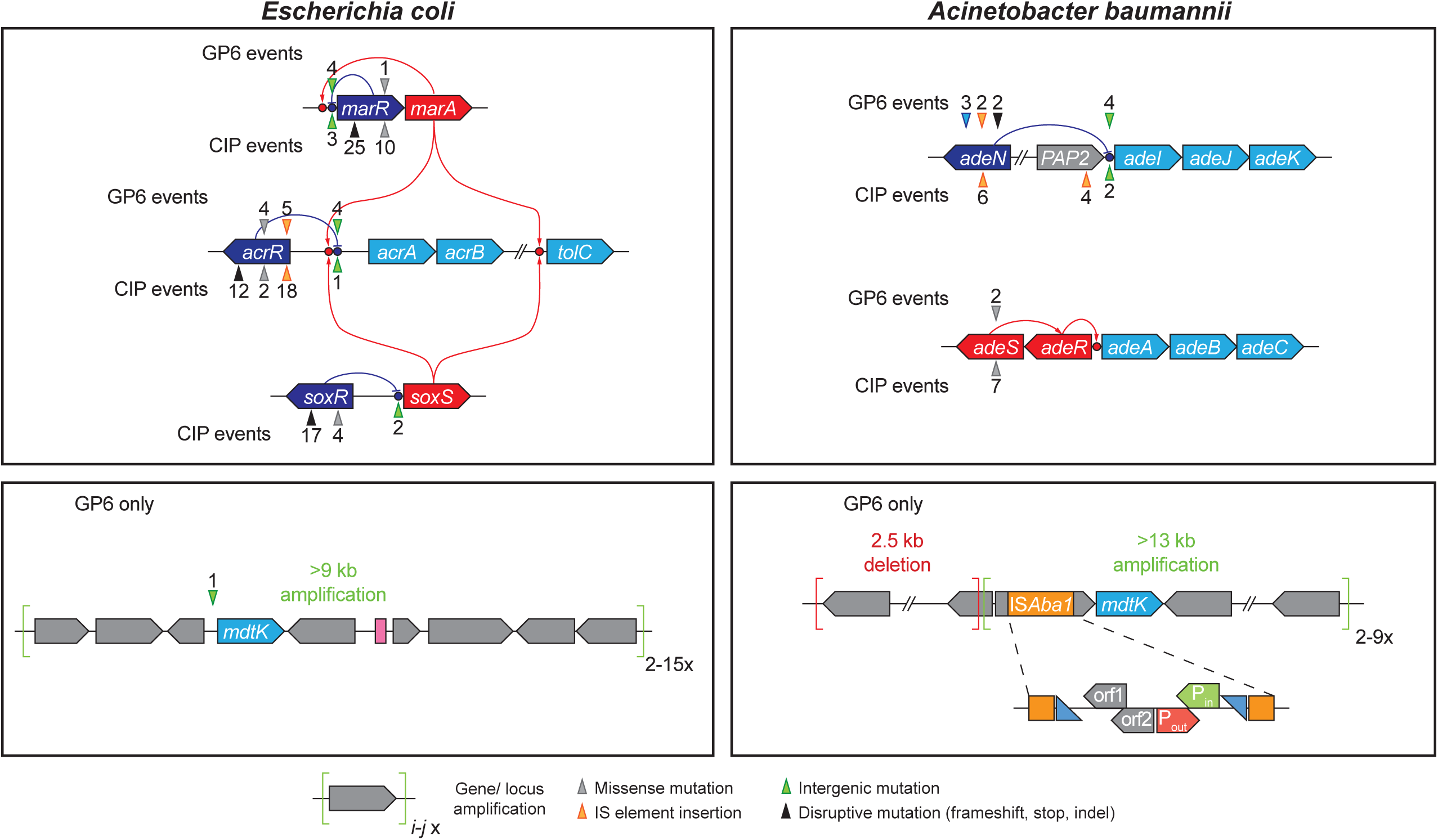
Mutational events in efflux systems observed in experimental evolution of GP6 resistance in *E. coli* and *A. baumannii*; comparison with CIP resistance studies ^14^. Gene coloring scheme: light blue – components of efflux transporters; dark blue – transcriptional repressors; red – transcriptional activators (including two-component systems); Grey – other genes. GP6-selected mutational events are shown above the affected genes, CIP-selected – below the genes. The positions of mutational events are shown by triangles (colored depending on type of events as shown below the diagram) and numbers reflecting the number of their independent occurrences.

Remarkably, the amplification of multi-gene genomic loci containing the *mdtK* gene, encoding a MATE family transporter implicated in efflux of multiple drugs including quinolones ^21^, massively occurred in both species during the evolution of resistance to GP6, but not CIP **(Table 1)**. The CNV analysis of *E. coli* WGS data showed that this locus was amplified from 2x to 17x in 5 out of 6 reactors (see **Supplementary Table S1A**). In addition, a short (50 bp) deletion was observed in the intergenic region upstream of the *mdtK* gene also leading to its overexpression (as further confirmed by clonal analysis, **Supplementary Figure S4**) via an unknown mechanism. In *A. baumannii* we observed three distinct mechanisms, all potentially contributing to *mdtK* overexpression: (i) 3-11x amplification of ≥13 kb genomic locus containing the *mdtK* gene in 5 out of 6 reactors; (ii) insertion of the *ISAba1* element providing a duly oriented strong promoter ^22^ immediately upstream of *mdtK* gene in all 6 reactors and reaching A_max_∼100% in 5 of them; and (iii) upstream deletion of a ∼2.5 kb genomic locus placing it under control of a promoter driving strong constitutive transcription of tRNA^Lys^(TTT)-encoding gene (**Figure 3 and Supplementary Table S1B**). These genomic rearrangements and their impact on *mdtK* overexpression were confirmed and refined by Nanopore sequencing and RT-qPCR analyses of selected clones (**Supplementary Figure S4**). Notably, *mdtK* genes of *E. coli* and *A. baumannii* comprise the only pair of orthologs shared by the respective multigene loci in the two species. This observation provided the initial implication of MdtK efflux transporter as a likely driver of GP6 resistance.

Other mechanisms of efflux deregulation in *A. baumannii* are largely overlapping between GP6 and CIP resistance studies (**Table 1** and **Figure 3**). Indeed, the most prominent of these in the evolution of GP6 resistance were two classes of events associated with the regulation of the AdeIJK efflux pump ^23^: (i) disruptive mutations (stop-gain, frameshift, and IS insertions) in the *adeN* gene encoding a transcriptional repressor of the *adeIJK* operon ^24^; and (ii) *ISAba1* insertions upstream of *adeIJK* operon providing a strong promoter likely leading to its overexpression. In one of the reactors, we observed two missense variants of the AdeS histidine kinase from the AdeRS two-component system, a known transcriptional activator of the AdeABC efflux pump ^25^. Such variants, far more prominent in the CIP study ^14^, have been implicated as drivers of multidrug resistance in *A. baumannii* ^25–28^.

As already emphasized, the main differences observed between evolutionary trajectories toward GP6 and CIP resistance were in the dynamics and frequency of efflux-driven vs target-driven mutational variants (**Figure 2**). Indeed, the evolution of CIP resistance in both species was invariably dominated by target modifications that emerged at the earliest stage and sustained at 100% abundance coupling with secondary efflux-upregulating events at the later stage. In contrast, target modifications were relatively scarce and even optional in the evolution of GP6 resistance in both, *E. coli* and *A. baumannii*. In the latter species, these variants emerged only at the later stage of evolution in addition to efflux-upregulating mutations (see **Figure 4 and Supplementary Figure S5**).

**Figure 4.**
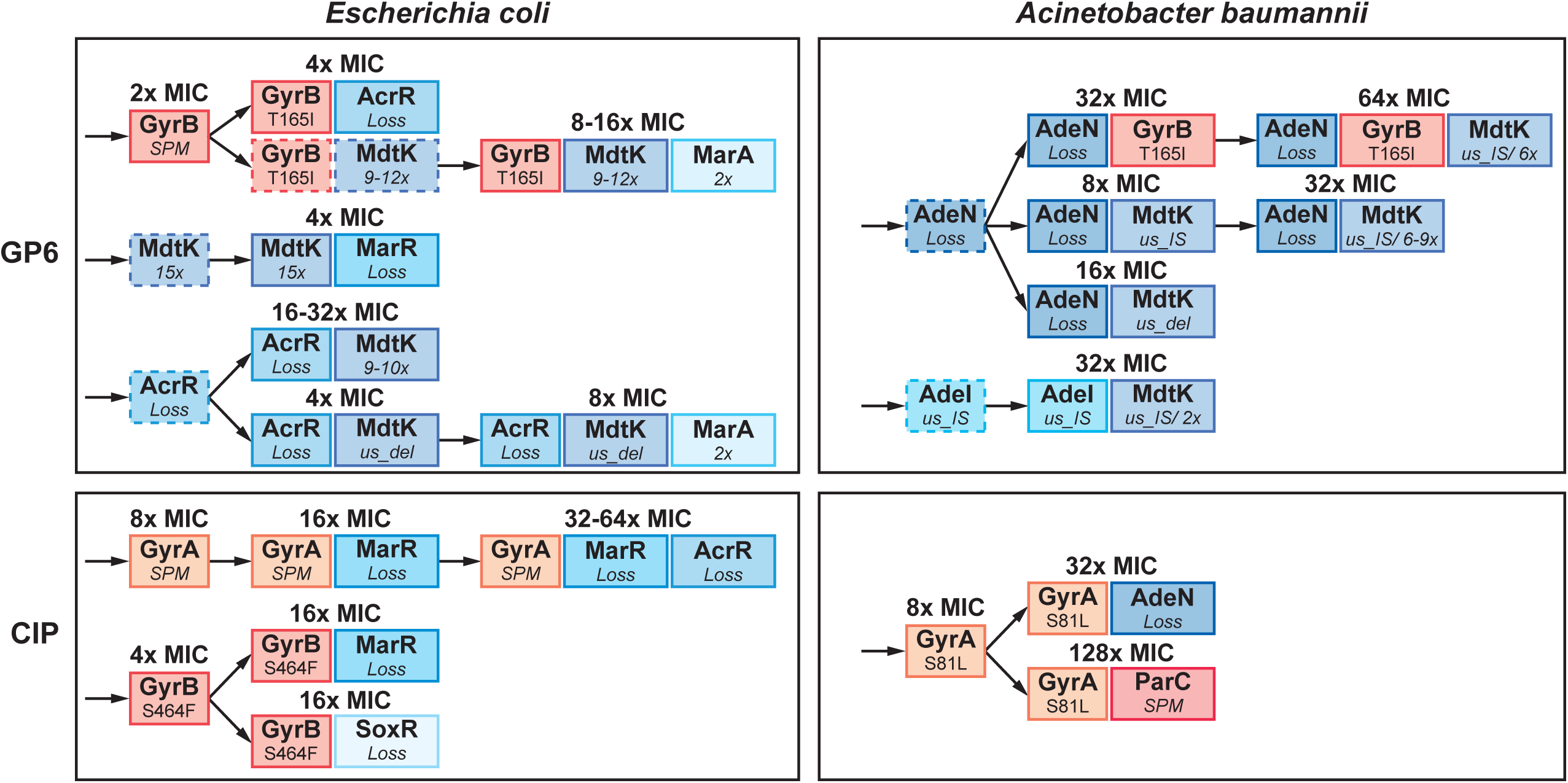
The order of occurrence and impact of mutations and their combinations driving the evolution of resistance to GP6 in *E. coli* and *A. baumannii*, a comparison with the experimental evolution of CIP resistance (modified from (14)). Only main driver mutations represented in characterized clones (from **Supplementary Table S3ABCD**) are shown leading to: (i) target modification (orange background palette); and (ii) efflux upregulation (blue background palette). The impact of these mutations on MIC (fold change) of respective drugs (MIC^GP6^ or MIC^CIP^) is shown above the corresponding variants. Intermediate variants (mostly, single mutants not represented in isolated clones) hypothesized based on dynamic plots (Supplementary Figure S3) are indicated by dashed borders and arrows.

### Other potentially significant variants

Among the 4 genes implicated by potentially significant mutational variants observed during experimental evolution of GP6 resistance in *E. coli,* the most prominent is a global transcriptional regulator Rob featuring three amino acid substitutions in close spatial proximity (**Supplementary Figure S5**). These features are consistent with a functional modification rather than with a loss of function. Although this protein was tentatively implicated in antibiotic resistance ^29,30^, the impact of the observed amino acid substitutions in GP6 resistance or compensatory fitness (if any) remains unclear.

Three additional *E. coli* genes, *rpoC, lon* and *yadG*, were implicated by a single mutational variant each but deemed potentially significant as they reached high A_max_ (in a range of 75 – 97%) and sustained until the end of the evolutionary experiment (see comments to **Supplementary Figure S3AB** for further details). Among them, the RpoC: Gly336Ser variant affecting the β’-subunit of the DNA-directed RNA polymerase complex (RNAP) was previously reported in clinical isolates of pathogenic *E. coli* causing urinary tract infections ^31,32^. An insertion of the *IS186B* mobile element upstream of the *lon* gene, a known hotspot for *IS186* insertion, is expected to downregulate Lon protease activity ^33^ leading to stabilization of SoxS and MarA, positive regulators of the AcrAB/TolC efflux system ^34,35^ and thus to upregulation of efflux. A single frameshift variant YadG:Val83fs would lead to a functional loss of the ATPase component of an uncharacterized ABC-type transporter of the Drug Exporter-1 family ^36^, the functional significance of which is unclear.

Three of the four *A. baumannii* genes tentatively implicated in evolution of GP6 resistance, *ettA, rplA and mnmG,* are associated with translation machinery. Two of them feature variants leading to a loss-of-function of: (i) the EttA protein which controls ATP/ADP ratio-dependent ribosomal entry into translation elongation phase ^37^; and (ii) the RplA ribosomal protein L1. In-frame insertion modifying the MnmG enzyme involved in posttranscriptional modification of tRNA, generating 5-methylaminomethyl-2-thiouridine in the wobble position 34 ^38,39^, was observed in 4 reactors. Functional significance of these events (if any) remains unclear. Observed mutational variants of RpoB, the β-subunit of RNAP, are of potential interest since some mutations in *rpoB* were previously implicated with CIP resistance and upregulation of the *mdtK* gene ^32^.

Overall, there is no direct evidence supporting contribution of these mutational variants to GP6 resistance or compensatory fitness, and none of the respective genes were previously implicated in evolution of CIP resistance ^14^.

### Clonal analysis and GP6/CIP cross-resistance

We used the bacterial population deconvolution algorithm to optimize a clone selection strategy from the samples collected during experimental evolution of GP6 resistance in both species (as described in **Supplementary Methods**). This approach facilitates the rational selection of the samples from which to isolate clones and determines the optimal number of clones for isolation to maximize representation of major mutational variants while minimizing redundancy. The complete results of WGS and MIC^GP6^ measurements for two sets of nonredundant clones, 12 from *E. coli* and 11 from *A. baumannii*, are provided in the **Supplementary Table S2AB**.

#### Genotype-to-phenotype mapping in selected clones from the experimental evolution of GP6 resistance in E. coli and A. baumannii

The accomplished clonal analysis yielded direct verification of major mutational events and their combinations in double and triple mutants. Among the *E. coli* clones, we observed: (i) both major target modifications (GyrB:Thr165Ile and GyrB:Gly77Ser) in 6 nonredundant clones; (ii) several mutational events associated with upregulation of AcrAB/TolC efflux systems including major missense mutations and IS-insertions in AcrR (8 clones) and MarR (1 clone) transcriptional regulators; (iii) 2x amplification of the *marR-marAB* divergon (4 clones); (iv) intergenic mutations upstream of the MdtK efflux transporter encoding gene (3 clones) and 9-15x amplification of a large (∼ 20 kB) *mdtK* containing genomic locus (5 clones); and (v) mutational variants affecting 3 out of 4 other potentially relevant genes *lon, rpoC* and *rob*, each represented by a single clone (**Supplementary Table S2A**).

A set of characterized *A. baumannii* clones featured: (i) a single major GyrB:Thr179Ile variant (in 4 clones); (ii) 3 variants affecting regulation of the AdeIJK efflux pump including two disruptive events in the AdeN repressor and a mutation in its likely operator site upstream of the *adeIJK* operon (collectively spanning all 11 clones); (iii) all three types of events that should lead to overexpression of the MdtK efflux transporter: upstream IS insertion (7 clones); large deletion (∼3.5 kb) of an upstream locus (3 clones), and 2-9x amplification of a large (∼ 12 kb) *mdtK*-containing genomic locus (4 clones); and (iv) single clones representing potentially significant mutational variants of 3 out of 4 tentatively implicated “other” genes, *rpoB, ettA,* and *mnmG* (**Supplementary Table S2B**).

Genomic rearrangements associated with the *mdtK* gene in both species (**Supplementary Figure S4A, B**) were directly confirmed by long-read Nanopore sequencing for representative clones of each species. An inferred overexpression of the *mdtK* gene resulting from these rearrangements was validated in both species by RT-qPCR in selected clones representing all types of mutational events (amplification, upstream deletion and IS-insertion) in this locus (**Supplementary Figure S4C, D**). Thus, in *A. baumannii*, IS-insertion upstream of *mdtK* gene led to ∼4-fold increase in its expression (relative to WT), while a combination of IS-insertion with 9x locus amplification further increased *mdtK* gene expression up to ∼64-fold. Interestingly, a ∼ 3.5 kb upstream deletion led to a comparable ∼45-fold increase in *mdtK* gene expression. Similar effects were observed in *E. coli* clones where the strongest (>20-fold) overexpression resulted from a 50-bp upstream deletion while a 15x locus amplification led to a more modest (∼7-fold) increase in *mdtK* expression.

Notably, isolated GP6^R^ clones of both species featured mostly double and triple mutants, which prevents accurate assessment of the impact of individual mutations on resistance or compensatory fitness. Nevertheless, from the observed 2-fold MIC^GP6^ increase in two *E. coli* clones featuring GyrB:Thr165Ile and GyrB:Gly77Ser/RpoC:Gly336Ser variants without any efflux-related mutations (see **Supplementary Table S3A**), we can deduce that these target modifications have a relatively small impact on GP6 resistance. This is in contrast with 8-fold MIC^CIP^ increase observed in single mutant variants GyrA:Asp87Tyr and GyrA:Ser83Leu from the CIP study ^14^ (**Supplementary Table S3B**). Each of the 6 characterized GP6-evolved *E. coli* clones without GyrB target modifications harbored at least two efflux-related mutations increasing their MIC^GP6^ in a range of 4–32x as compared to unevolved parental strain. Notably, each of these clones featured an event affecting MdtK expression. The remaining 4 clones combining the GyrB:Thr165Ile variant with 1-3 efflux-deregulation events displayed 4-16x increase of MIC^GP6^.

Dissection of MIC^GP6^ effects of individual variants was even more difficult for isolated *A. baumannii* clones which typically harbored 3-4 mutations per clone (**Supplementary Table S3C**). All 4 clones featuring the main target modification variant (GyrB:Thr179Ile) also featured IS inserts in the *adeN* gene and some additional events. Indirectly, we can estimate that the contribution of target modification to MIC^GP6^ increase is also relatively modest by a comparison of two nearly isogenic clones, 3F10 (GyrB:Thr179Ile/AdeN:IS/*us_mdtK*:IS/MdtK:6x) and 3D2 (AdeN:IS/*us_mdtK*:IS/MdtK:9x), displaying only a 2-fold difference in their MIC^GP6^ values (64x vs 32x relative to WT). Both efflux pumps, AdeIJK and MdtK, contribute strongly to GP6 resistance in *A. baumannii*, albeit their relative impacts cannot be confidently dissected. A cumulative effect of these two types of events is apparent from the comparison of clones: 1F1 (*adeN:IS*/*us_mdtK:IS*) and 2D2 (*adeN:IS/ us_mdtK:IS/*MdtK:5x), where MdtK amplification led to an additional 4-fold increase of MIC^GP6^. Although “other” mutations were captured in 1-2 clones (in combination with driver mutations), ascertaining their potential relevance would require additional experiments.

Evolutionary trajectories deduced from mutational mapping in both GP6^R^ and CIP^R^ clones (**Figure 5**) illustrate an already noted fundamental difference in the dynamics of resistance acquisition between the two drugs. Indeed, efflux-upregulating events start in the early stage and dominate the evolution of GP6 resistance in both species, while GyrB target modifications appear optional and emerging after efflux mutations (as in *A. baumannii*). This is in contrast with the previously observed 2-stage evolution of CIP resistance where GyrAB target modifications always preceded efflux-upregulating events ^14^.

**Figure 5.**
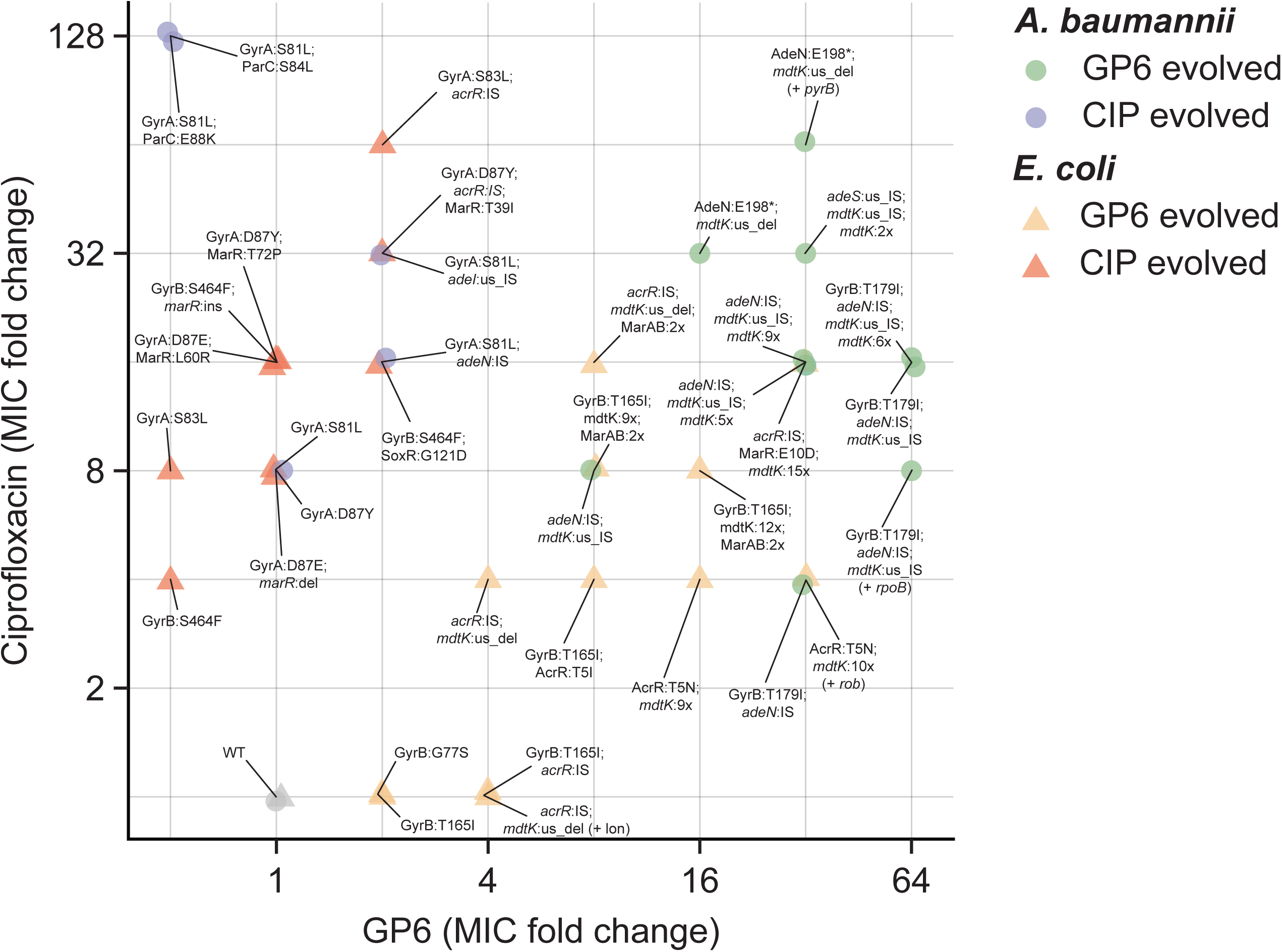
Correlation plot of changes in MIC values of the two compounds, GP6 and CIP. observed over a panel of ∼40 nonredundant clones selected from morbidostat-based studies on experimental evolution of resistance to GP6 (yellow and green symbols) or CIP (orange and light-brown symbols) in *E. coli* (triangles) and *A. baumannii* (circles). Corresponding mutational variants (only driver mutations) are shown as data labels using a matching font color. Complete mutational and MIC data are provided in **Supplementary Table S3ABCD**. Plotted values reflect fold-change of MIC (FC) measured for both drugs in each clone as compared to a respective MIC of the unevolved parental strain. Two outlined areas illustrate a clear separation between CIP^R^-evolved isolates (that do not show any increase in MIC to GP6) and GP6 ^R^-evolved isolates (many of which show an appreciable cross-resistance to CIP).

#### GP6 vs CIP cross-resistance analysis of E. coli and A. baumannii clones isolated from experimental evolution to GP6 or CIP

To further elucidate shared and unique resistance drivers for the two pharmacologically distinct drugs, we measured orthogonal MIC^CIP^ values for all clones selected in GP6 experimental evolution of both species. Likewise, MIC^GP6^ values were measured for representative clones selected from CIP experimental evolution ^14^. A comparison of the obtained cross-resistance data is provided in **Supplementary Table S3 and Figure 4**.

A global comparison of MIC effects as reflected in a correlation plot (**Figure 5**) reveals one obvious trend. While GP6-evolved mutational variants show a clear tendency to display strong CIP resistance, no such cross-resistance is observed for CIP-evolved clones. Indeed, only 4 clones (all in *E. coli*) out of 23 analyzed GP6-evolved clones (from both species) displayed unchanged MIC^CIP^ as compared to respective unevolved strains, while a 4-64x increase in MIC^CIP^ was observed for all other GP6-evolved clones. Notably, 2 of the 4 CIP-susceptible clones are driven by GyrB variants without any efflux-related events. This is consistent with the obviously nonoverlapping distribution of target-based mutations driving GP6 vs CIP resistance (see **Figure 2**) and with the lack of any impact of CIP-evolved target modifications on MIC^GP6^ (**Supplementary Table S3B, D**).

As already discussed, most GP6-evolved *E. coli* clones and all but one *A. baumannii* clone harbor at least two different efflux-related mutational events, which complicates the dissection of their individual impacts on MIC for either drug. Nevertheless, there is no doubt that a combination of two or more mutational events in efflux-driving genes emerging in both species under evolutionary pressure of GP6 yielded a robust CIP resistance without any contribution by target modifications. This observation is in contrast with our previous study ^14^ where the observed evolutionary trajectories to CIP resistance (**Figure 4**) and all of the isolated CIP^R^ clones included target modification variants.

One possible interpretation is that a ∼2x increase in MIC^GP6^ driven by mutations in the GP6-binding site of GyrB alone is insufficient for robust selection as compared to ∼8x increase of MIC^CIP^ driven by mutations in the CIP-binding site of GyrA/GyrB complex. A major difference between the evolutionary trajectories characteristic of these two drugs may be at least partially due to their different relative affinities for two targets, GyrA/GyrB and ParC/ParE. Indeed this is consistent with prior work, with suggestions that the greater affinity of CIP for GyrA (versus ParC) enables point mutations to occur only in this gene with a moderate increase in MIC^CIP^ ^40^, whilst GP6 has comparable affinities for both GyrB and ParE (though still a slightly greater affinity for GyrB) minimizing the impact of a single point mutation in just one of these topoisomerase complexes ^11^.

Both major *E. coli* GyrB variants observed in our study (GyrB:Gly77Ser and GyrB:Thr165Asn) are known as drivers of novobiocin resistance decreasing ATPase and DNA supercoiling activities of GyrAB ^41^. They may be associated with potentially higher fitness costs as compared to CIP^R^ GyrA variants known to have a negligible effect on fitness ^42^. Some or all of these considerations may explain why efflux upregulation, which is an optional/late event in evolution of CIP resistance, may turn into a mandatory/early mechanism on the path to GP6 resistance. Not surprisingly, single efflux-related events that are most common among CIP-evolved clones do not provide sufficient protection against GP6 as indicated by double or even multiple efflux upregulation variants in GP6-evolved clones (**Supplementary Table S3A, C**)

Another interesting question is why upregulation of the MdtK transporter was not observed in our previous study on experimental evolution of CIP resistance ^14^? Indeed, MdtK was previously reported to contribute to efflux of fluoroquinolones ^21,43,44^, which is further supported by our GP6-CIP cross-resistance analysis. A likely explanation is based on the same premise of a stronger requirement of efflux upregulation for establishing a robust GP6 vs CIP resistance. It is reasonable to surmise that upregulation of AcrAB/TolC (in *E. coli*) or AdeIJK (in *A. baumannii*) efflux pumps via disruptive regulatory mutations (**Figure 3**) may occur at higher frequency than overexpression of MdtK, which requires larger genomic rearrangements.

## Conclusions

Through a morbidostat-based experimental evolution approach, resistance to the TriBE inhibitor GP6 ^12^ was investigated in *E. coli* and *A. baumannii*. A comprehensive two-dimensional comparison of the mutational landscapes and dynamics was then performed, comparing these results with a previous study on the evolution of CIP resistance in the same species ^14^. This analysis yielded significant insights into the development of resistance towards distinct classes of bacterial type-II topoisomerase inhibitors.

In both species, the main drivers of GP6 resistance were either GyrB ATP-binding site mutations consistent with the mechanism of GP6 binding, or efflux upregulation events. The order in which these events were detected here was inverted relative to our previous CIP study (14), which is attributed to the comparable affinity of GP6 to both targets, GyrB and ParE preventing a meaningful increase in MIC^GP6^ with any single target mutation. That said, the accurate interpretation of the observed evolutionary trajectories would require the additional knowledge of competitive fitness of the emerging mutational variants.

While some mutational events affecting species-specific efflux systems (AcrAB/TolC in *E. coli* and AdeIJK/AdeAB in *A. baumannii*) are shared between GP6 and CIP experimental evolution, the upregulation of the MdtK efflux pump in both species was prominent and impactful only during the evolution of resistance to GP6.

From cross-resistance experiments, it was observed that most isolated GP6 evolved clones were also resistant to CIP whilst the opposite was not true. Despite this potential contribution of the MdtK-driven efflux to CIP resistance of GP6-evolved clones, no mutational variants affecting MdtK were detected during evolution of CIP resistance. This apparent disparity may reflect a lower probability of genomic rearrangements (that are required for MdtK upregulation) as compared to disruptive mutations (that are sufficient to upregulate other efflux systems) given the diminished contributions of efflux under the target-driven evolution of CIP resistance. Overall, we have illustrated the utility of our morbidostat-based experimental evolution workflow for the elucidation of dynamics and mechanisms underlying acquisition of antibiotic resistance. Despite the obvious limitations of in vitro methodology, we believe that the employment of such a workflow is of utmost importance for the ability to infer a resistibility potential of bacterial pathogens toward various antibiotics including newly developed compounds and those in development. Additionally, we expect this to become a tool used to aid the rational selection of drug(s), aiming to minimize the selection of ineffective treatment regimens. For example, both this study on GP6 and previous research on CIP suggest that GP6 treatment would effectively target bacterial pathogens with acquired CIP resistance, whereas the reverse is not the case.

## Acknowledgments

This work was supported by F. Hoffmann-La Roche Ltd and by NIAID grant 5R01AI167977 to A.L.O.

## Author Contributions

S.A.L. planned and oversaw morbidostat experiments, performed sequencing data processing, bioinformatics analysis and drafting the manuscript. J.E.K. performed RT-qPCR experiments and contributed to manuscript writing and editing. J.E.Z. performed morbidostat experiments and clonal analysis, contributed to manuscript editing. M.L.E. assisted in the performance of morbidostat experiments, prepared genomic libraries for sequencing and measured MIC values. M.V. provided synthetic compounds and participated in general planning, discussion, and MS editing. A.L.O. provided general project planning, oversight, data interpretation, led in manuscript writing and editing. Semen A. Leyn and James E. Kent contributed equally to this work.

## SUPPLEMENTARY MATERIALS

**A. Supplementary Methods** (file GyrB_GP6_supplementary_methods_Draft4.docx).

- MORBIDOSTAT SETUP AND PROGRAMMING

- SEQUENCING DATA ANALYSIS, VARIANT CALLING AND RANKING

POPULATION WGS DATA DECONVOLUTION FOR EVOLUTIONARY DYNAMICS AND CLONAL ANALYSIS RNA ISOLATION AND RT-QPCR

**B. Supplementary Tables:**

**Supplementary Table S1.** WGS data for evolving bacterial populations. (File “GyrB_GP6_Supplementary table S1ABC_Draft4.xlsx”)

**Table S1A.** Relative abundances (%) of mutational variants and CNV numbers observed in sequencing data obtained for population samples (in time series) of *E. coli* BW25113 from morbidostat-based experimental evolution of resistance to GP6 compound. (For CNV analysis only genes with observed values ≥ 2 or ≤0.5 and SD values among samples ≥ 0.1 are reported.)

**Table S1B.** Relative abundances (%) of mutational variants and CNV numbers observed in sequencing data obtained for population samples (in time series) of *A. baumannii* ATCC17978 from morbidostat-based experimental evolution of resistance to GP6 compound.

**Table S1C.** Quality metrics for short read alignments of populations samples.

**Supplementary Table S2.** WGS and MIC data for selected clones.

(File “GyrB_GP6_Supplementary table S2ABC_Draft4.xlsx”)

**Table S2A.** Mutational variants observed in sequencing data and measured MIC values of selected clones from morbidostat-based experimental evolution of resistance to GP6 compound in *E. coli* BW25113. (For CNV analysis only genes with observed values ≥ 2 or ≤0.5 and SD values among samples ≥ 0.1 are reported.)

**Table S2B.** Mutational variants observed in sequencing data and measured MIC values of selected clones from morbidostat-based experimental evolution of resistance to GP6 compound in *A. baumannii* ATCC17978.

**Table S2C**. Quality metrics for short read alignments of clone samples.

**Supplementary Table S3. ABCD.** Genotype-phenotype associations in representative clones selected from the experimental evolution of resistance to GP6 (A, C) in *E. coli* and comparison with CIP^R^ clones (B, D) in *E. coli* (A, B) and *A. baumannii* (C, D).

(File “GyrB_GP6_Supplementary_Table_S3_Draft4.docx”)

**Supplementary Table S4. AB.** WGS data for bacterial populations of *E. coli* and *A. baumannii* upon non-selective continuous culturing

**Table S4A.** Relative abundances (%) of mutational variants observed in sequencing data obtained for *E. coli* BW25113 from non-selective continuous culturing in turbidostat mode.

**Table S4B.** Relative abundances (%) of mutational variants observed in sequencing data obtained for *A. baumannii* ATCC17978 from non-selective continuous culturing in turbidostat mode.

**Table S4C.** Coverage for short read alignments.

**C. Supplementary Figures** (File GyrB_GP6_Supplementary_figures_revised_1.docx)

**Supplementary Figure S1.** General workflow diagram of morbidostat-based evolution of resistance to GP6 compound in *E. coli* BW25113 and *A. baumannii* ATCC17978.

**Supplementary figure S2.** OD profiles (black line) and calculated drug concentration (red line) profiles for morbidostat-based experimental evolution of resistance to GP6 in *E. coli (S2A) and A. baumannii (S2B*). Collected samples are shown by arrows and labeled with a time stamp of collection.

**Supplementary Figure S3.** Dynamics of major mutational variants in morbidostat-based evolution of resistance to GP6 compound in *E. coli* BW25113 (A) and *A. baumannii* ATCC17978 (B).

Cumulative area plots are shown for the most prominent mutational variants reaching relative abundance Amax ≥10% in at least one sample. Plots are shown for each of the six reactors R1-R6. Y-axis shows variant abundance (%) in bacterial population samples taken at corresponding time points (X-axis). Red line marks a 100% threshold, which helps to visualize the emergence of double and triple mutants. *Gene names and corresponding functions are explained in the legend to Table 1 (and in Supplementary Table S1A with more details and tabular data)*.

**Supplementary Figure S4.** Examples of genomic rearrangement in the loci harboring *mdtK* multidrug efflux transporter gene in *E. coli* (A) and *A. baumannii* (B, C). The details of Nanopore-based verification and mapping of genomic amplification are illustrated for clones: 4F9 (*E. coli*) and 3D2 (*A. baumannii*)

**Supplementary Figure S5.** Structure of Rob transcriptional regulator complex with DNA (PDB:1D5Y ^45^). Residues with variants observed in GP6 morbidostat experiments are colored red.

