## Supplemental table 3 for "Two Classes of DNA Gyrase Inhibitors Elicit Distinct Evolutionary Trajectories Toward Resistance in Gram-Negative Pathogens"

**Supplementary Table S3.** Genotype to phenotype associations in representative clones selected from the experimental evolution of resistance to GP6 in *E. coli* (A) and comparison with CIP-resistant clones (B) from (3).

**A**

| ***E.coli*, GP6-evolved** | | WT | 1F3 | 5B2 | 5C1 | 5F1 | 5F2 | 5C2 | 2F4 | 2F3 | 2F1 | 4F9 | 6F7 | 6F6 |
| --- | --- | --- | --- | --- | --- | --- | --- | --- | --- | --- | --- | --- | --- | --- |
| GyrB:Gly77Ser | **T** | - | + | - | - | - | - | - | - | - | - | - | - | - |
| GyrB:Thr165Ile | **T** | - | - | + | + | + | + | + | - | - | - | - | - | - |
| *acrR:IS* | **E** | - | - | - | + | - | - | - | + | + | + | + | - | - |
| AcrR:Thr5Ile | **E** | - | - | - | - | - | - | + | - | - | - | - | - | - |
| AcrR:Thr5Asn | **E** | - | - | - | - | - | - | - | - | - | - | - | + | + |
| MarR:Glu10Asp | **E** | - | - | - | - | - | - | - | - | - | - | + | - | - |
| *us_mdtK*:del | **E** | - | - | - | - | - | - | - | + | + | + | - | - | + |
| *mdtK*:amp (x) | **E** | 1 | 1 | 1 | 1 | **9** | **12** | 1 | 1 | 1 | 1 | **15** | **9** | **10** |
| *marRAB*:amp (x) | **E** | 1 | 1 | 1 | 1 | **2** | **2** | 1 | 1 | 1 | **2** | 1 | 1 | 1 |
| Rob:Cys147Tyr | O | - | - | - | - | - | - | - | - | - | - | - | - | + |
| *us_lon:IS* | O | - | - | - | - | - | - | - | - | + | - | - | - | - |
| RpoC:Gly336Ser | O | - | + | - | - | - | - | - | - | - | - | - | - | - |
| YadG:Val83fs | O | - | - | - | - | + | + | - | - | - | - | - | - | - |
| MIC^GP6^, fold change vs WT | | **1** | **2** | **2** | **4** | **8** | **16** | **8** | **4** | **4** | **8** | **32** | **16** | **32** |
| MIC^CIP^, fold change vs WT | | **1** | **1** | **1** | **1** | **8** | **8** | **4** | **4** | **1** | **16** | **16** | **4** | **4** |

B

| ***E.coli,* CIP-evolved** | | WT | 6G64 | 2G23 | 2G17 | 5E1 | 1G3 | 3F3 | 4D2 | 4G39 | 4D1 | 5E2 | 5E4 |
| --- | --- | --- | --- | --- | --- | --- | --- | --- | --- | --- | --- | --- | --- |
| GyrA:Asp87Tyr | **T** | - | + | + | + | - | - | - | - | - | - | - | - |
| GyrA:Asp87Gly | **T** | - | - | - | - | + | + | + | - | - | - | - | - |
| GyrA:Ser83Leu | **T** | - | - | - | - | - | - | - | + | + | - | - | - |
| GyrB:Ser464Phe | **T** | - | - | - | - | - | - | - | - | - | + | + | + |
| *acrR:IS* | **E** | - | - | - | + | - | - | - | - | + | - | - | - |
| MarR:Thr39Ile | **E** | - | - | - | + | - | - | - | - | - | - | - | - |
| MarR:Leu60Arg | **E** | - | - | - | - | + | - | - | - | - | - | - | - |
| MarR:Thr72Pro | **E** | - | - | + | - | - | - | - | - | - | - | - | - |
| *marR:del* | **E** | - | - | - | - | - | + | + | - | + | - | - | - |
| *marR:ins* | **E** | - | - | - | - | - | - | - | - | - | - | + | - |
| SoxR:Gly121Asp | **E** | - | - | - | - | - | - | - | - | - | - | - | + |
| MIC^GP6^, fold change vs WT | | **1** | **1** | **1** | **2** | **1** | **1** | **1** | **0.5** | **2** | **0.5** | **1** | **2** |
| MIC^CIP^, fold change vs WT | | **1** | **8** | **16** | **32** | **16** | **8** | **8** | **8** | **64** | **4** | **16** | **16** |

**Supplementary Table S3 (*continued*).** Genotype to phenotype associations in representative clones selected from the experimental evolution of resistance to GP6 in *A. baumannii* (C) and comparison with CIP-resistant clones (D) from the previous study (3).

**C**

| ***A.baumannii,* GP6-evolved** | | WT | 1F2 | 2F2 | 2F1 | 3F10 | 1F1 | 2D2 | 3D2 | 4F10 | 4F8 | 4D4 | 5F3 |
| --- | --- | --- | --- | --- | --- | --- | --- | --- | --- | --- | --- | --- | --- |
| GyrB:Thr179Ile | **T** | - | + | + | + | + | - | - | - | - | - | - | - |
| *adeN:IS* | **E** | - | + | + | + | + | + | + | + | - | - | - | - |
| AdeN:Glu198* | **E** | - | - | - | - | - | - | - | - | + | + | + | - |
| *us_adeI:IS* | **E** | - | - | - | - | - | - | - | - | - | - | - | + |
| *us_mdtK:IS* | **E** | - | - | + | + | + | + | + | + | - | - | - | + |
| us mdtK:del | **E** | - | - | - | - | - | - | - | - | + | + | + | - |
| *mdtK*:amp | **E** | 1 | 1 | 1 | 1 | **6** | 1 | **5** | **9** | 1 | 1 | 1 | **2** |
| RpoB:Asn777_ins | **O** | - | - | - | + | - | - | - | - | - | - | - | - |
| *ettA*:IS | **O** | - | - | + | - | - | - | - | - | - | - | - | - |
| MnmG:Val11_Ile12dup | **O** | - | + | - | - | - | - | - | - | - | - | - | - |
| PyrB:Thr76Ile | **O** | - | - | - | - | - | - | - | - | + | + | - | - |
| pAB3 plasmid:loss | **O** | - | + | - | - | - | - | - | - | + | - | - | - |
| MIC^GP6^, fold change vs WT | | **1** | **32** | **64** | **64** | **64** | **8** | **32** | **32** | **32** | **32** | **16** | **32** |
| MIC^CIP^, fold change vs WT | | **1** | **4** | **16** | **8** | **16** | **8** | **16** | **16** | **64** | **64** | **32** | **32** |

**D**

| ***A.baumannii,* CIP-evolved** | | WT | 6E9 | 5E5 | 3E1 | 5E1 | 4E7 | 6E8 |
| --- | --- | --- | --- | --- | --- | --- | --- | --- |
| GyrA:Ser81Leu | **T** | - | + | + | + | + | + | + |
| ParC:Ser84Leu | **T** | - | - | - | - | + | + | - |
| ParC:Glu88Lys | **T** | - | - | - | - | - | - | + |
| *adeN:IS* | **E** | - | - | + | - | - | - | - |
| *us_adeI:IS* | **E** | - | - | - | + | - | - | - |
| *us_csuA/B:IS* | **O** | - | - | + | + | + | - | - |
| *us_csuA/B:SNP* | **O** | - | + | - | - | - | - | - |
| MIC^GP6^, fold change vs WT | | **1** | **1** | **2** | **2** | **0.5** | **0.5** | **0.5** |
| MIC^CIP^, fold change vs WT | | **1** | **8** | **16** | **32** | **128** | **128** | **128** |

MIC measurements in 4 nonredundant sets of clones selected from morbidostat-based experimental evolution of resistance vs GP6 compound in this study (A,C) and vs CIP (B,D) in E. coli (A,B) and A. baumannii (C,D). All clones were completely sequenced to identify mutational events (shown by "+") including locus amplification (shown by a numeric value of CNV). For comparative purposes, MIC values for each drug are shown as fold change over MIC of the respective unevolved parental strains (WT). For SNV events, genes are denoted as amino acid substitutions in respective protein products (eg GyrB:Gly77Ser). Other events including IS insertions (IS) and indels in coding or in upstream ("*us_*") noncoding regions refer to respective gene IDs (eg *adeN:IS*). All genes and mutational events therein are assigned to one of the three categories (as in Table 1): “T” – target modification; “E” – efflux upregulation and “O” – other genes of yet unclear relevance (if any).

Category "Target" (both species):

**GyrB**, DNA gyrase subunit B;

**GyrA**, DNA gyrase subunit A;

**ParC,** Topoisomerase IV subunit A

Category "Efflux" (*E. coli*):

**AcrA**, membrane protein component of AcrAB multidrug efflux pump;

**AcrR**, transcriptional regulator of *acrAB* operon;

**MarR**, transcritional repressor of the multiple antibiotic resistance (*mar*) operon;

**SoxR**, Redox-sensitive transcriptional activator;

**MdtK**, Multidrug efflux transporter of MATE family;

Category "Efflux" (*A. baumannii*):

**AdeI,** membrane protein component of AdeIJK efflux pump.

**AdeN**, transcriptional repressor of multidrug efflux pump AdeIJK;

**AdeS,** histidine kinase of the two-component AdeRS system, transcriptional activator of the *adeAB* efflux pump operon

Category "Other" (*E.coli*):

**Rob,** global transcriptional regulator

**Lon,** ATP-dependent serine protease mediating unfolding and degradation of abnormal; and short-lived proteins.

**RpoC,** β'-subunit of DNA-directed RNA polymerase complex (RNAP)

**YadG,** ATPase component of an uncharacterized ABC-type transporter

Category "Other" (*A. baumannii*)*:

**RpoB,** DNA-directed RNA polymerase beta subunit

**EttA***,* Energy-dependent translational throttle protein A;

**MnmG**. tRNA-5-carboxymethylaminomethyl-2-thiouridine(34) synthesis protein;

**PyrB,** Aspartate carbamoyltransferase

**CsuA/B,** major CSU pilus subunit

*The only clone (1F2) carrying AdeN:IS and no MdtK-related events on the background of GyrB:Thr179Ile variant displayed MIC^GP6^ comparable to other clones (2F2, 2F1) carrying additionally *us_mdtK:IS* (5 uM vs 10 uM). Unfortunately, this comparison does not allow us to speculate about a relatively stronger impact of AdeIJK pump due to distinct other mutations observed in these clones (MnmG:Val11_Ile12dup vs *ettA:IS* or RpoB:Asn777_ins). In addition, one missense variant affecting aspartate carbamoyltransferase gene PyrB:Thr76Ile), which was not identified as potentially significant in population WGS data was mapped in two selected *A. baumannii* clones on the AdeN:Glu198*/*us_mdtK:IS* background (clones 4F8 and 4F10). Both clones showed a 2x increase of both MIC^GP6^ and MIC^CIP^ compared to a clone 4D4 lacking this additional variant. However, despite some data pointing to a cross-talk between CIP treatment and pyrimidine biosynthesis (13, 45), in the absence of support from population data, the observed modest MIC effect does not provide sufficient evidence of PyrB:Thr76Ile variant functional significance.
