## Supplemental figures for "Two Classes of DNA Gyrase Inhibitors Elicit Distinct Evolutionary Trajectories Toward Resistance in Gram-Negative Pathogens"

### Supplementary Figures

#### Supplementary Figure S1.

General workflow diagram of morbidostat-based evolution of resistance to GP6 compound in *E. coli* BW25113 and *A. baumannii* ATCC17978

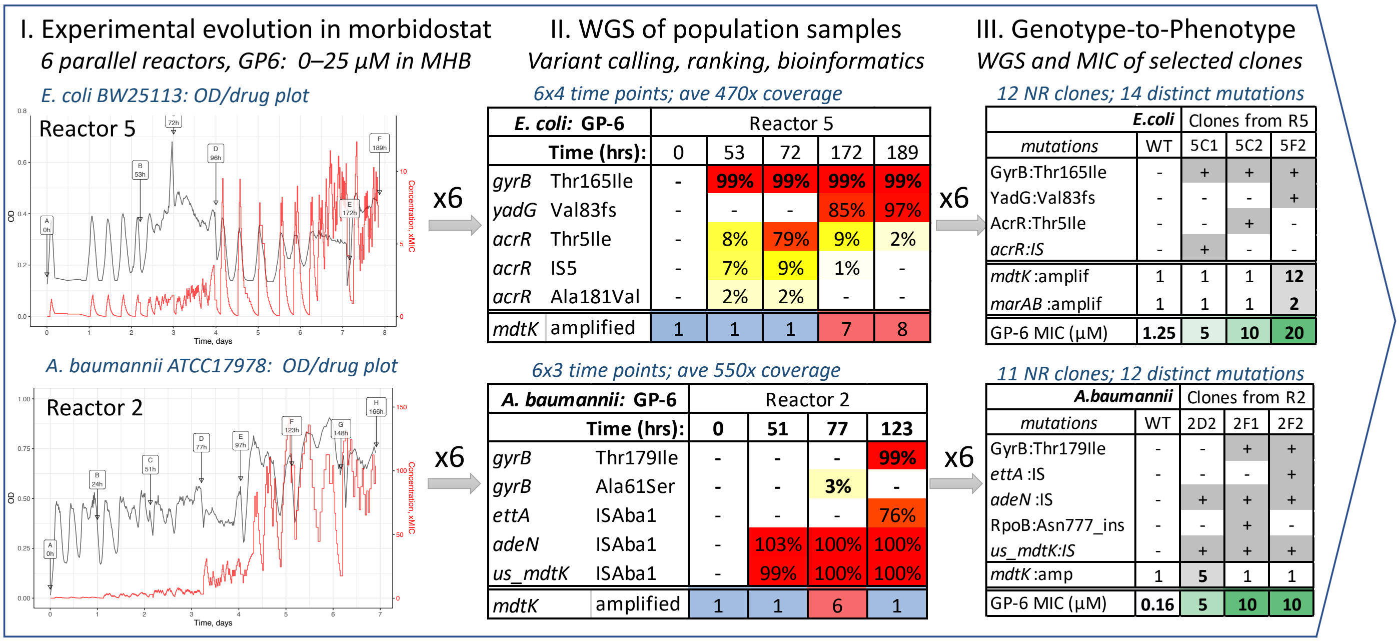

Supplementary figures

#### Supplementary figure S2.

**A:** OD profiles (black line) and calculated drug concentration (red line) profiles for experimental evolution in morbidostat of *E. coli*. Collected samples are shown by arrows and labeled with time.

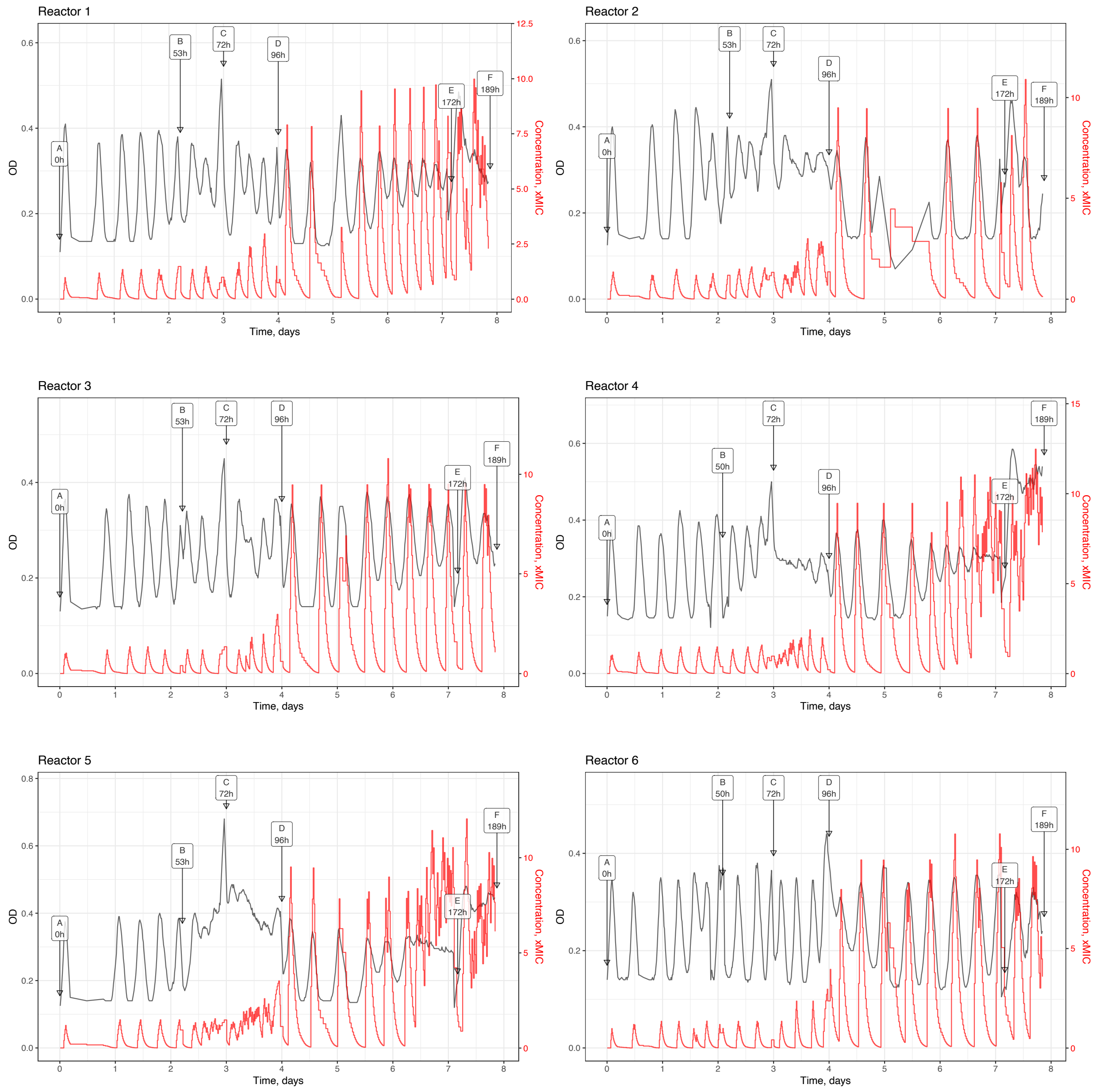

**Supplementary figure S2 (continued)**

**B:** OD profiles (black line) and calculated drug concentration (red line) profiles for experimental evolution in morbidostat of A. baumannii. Collected samples are shown by arrows and labeled with time.

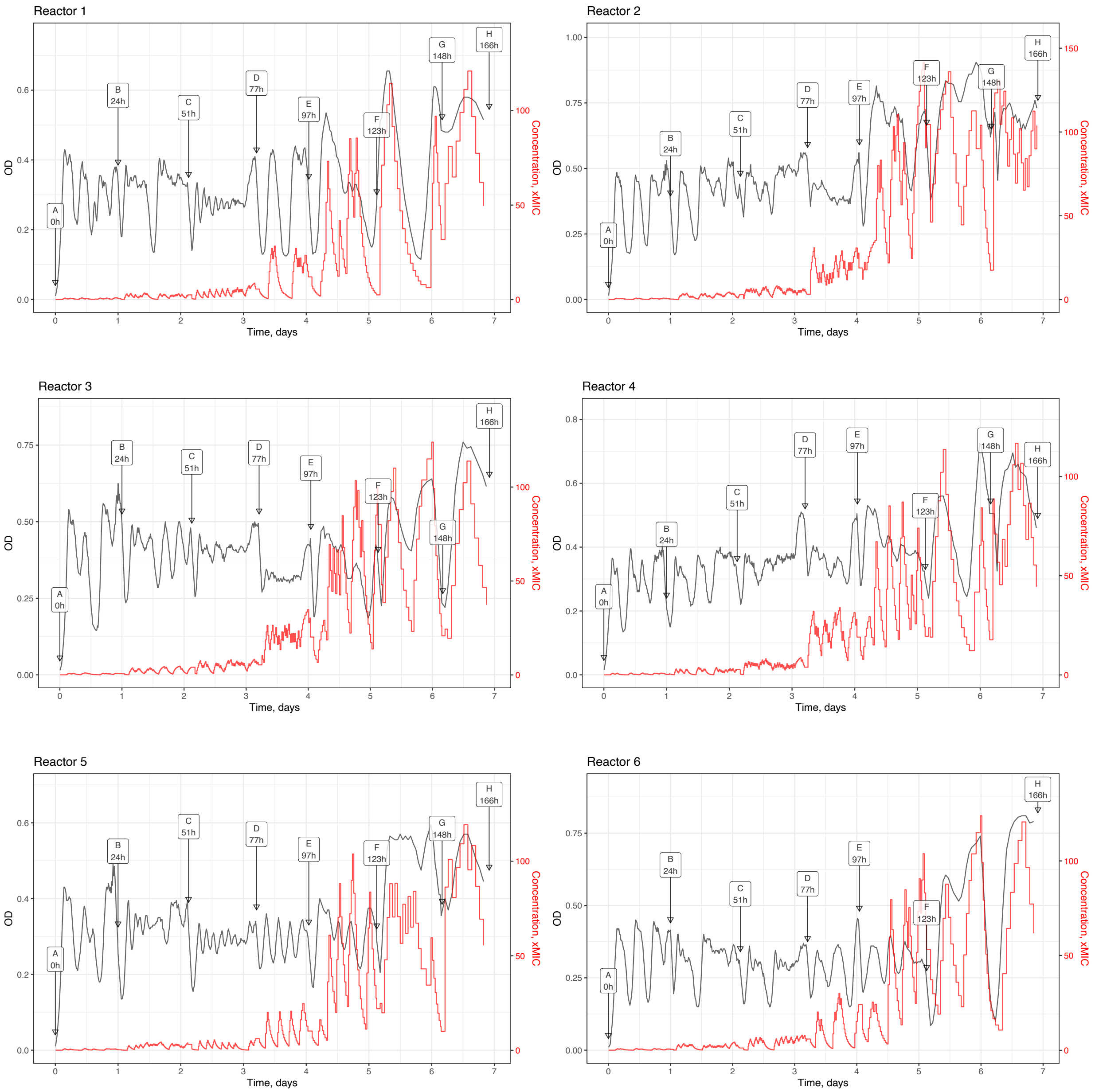

Supplementary figures

**Supplementary Figure S3.** Dynamics of major mutational variants in morbidostat-based evolution of resistance to GP6 compound in *E. coli* BW25113 (A) and *A. baumannii* ATCC17978 (B).

**Supplementary Figure S3A**

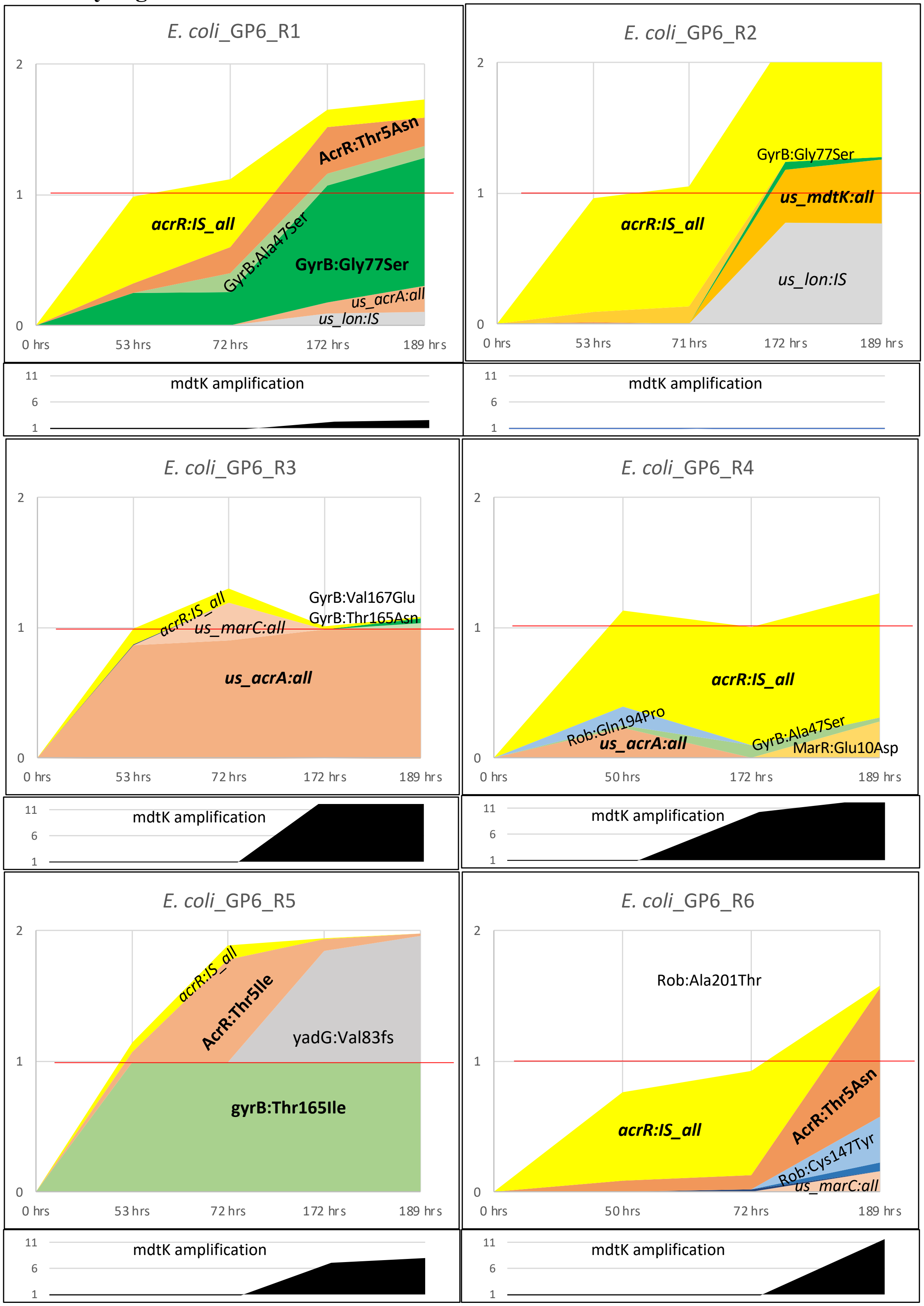

**Supplementary Figure S3B**

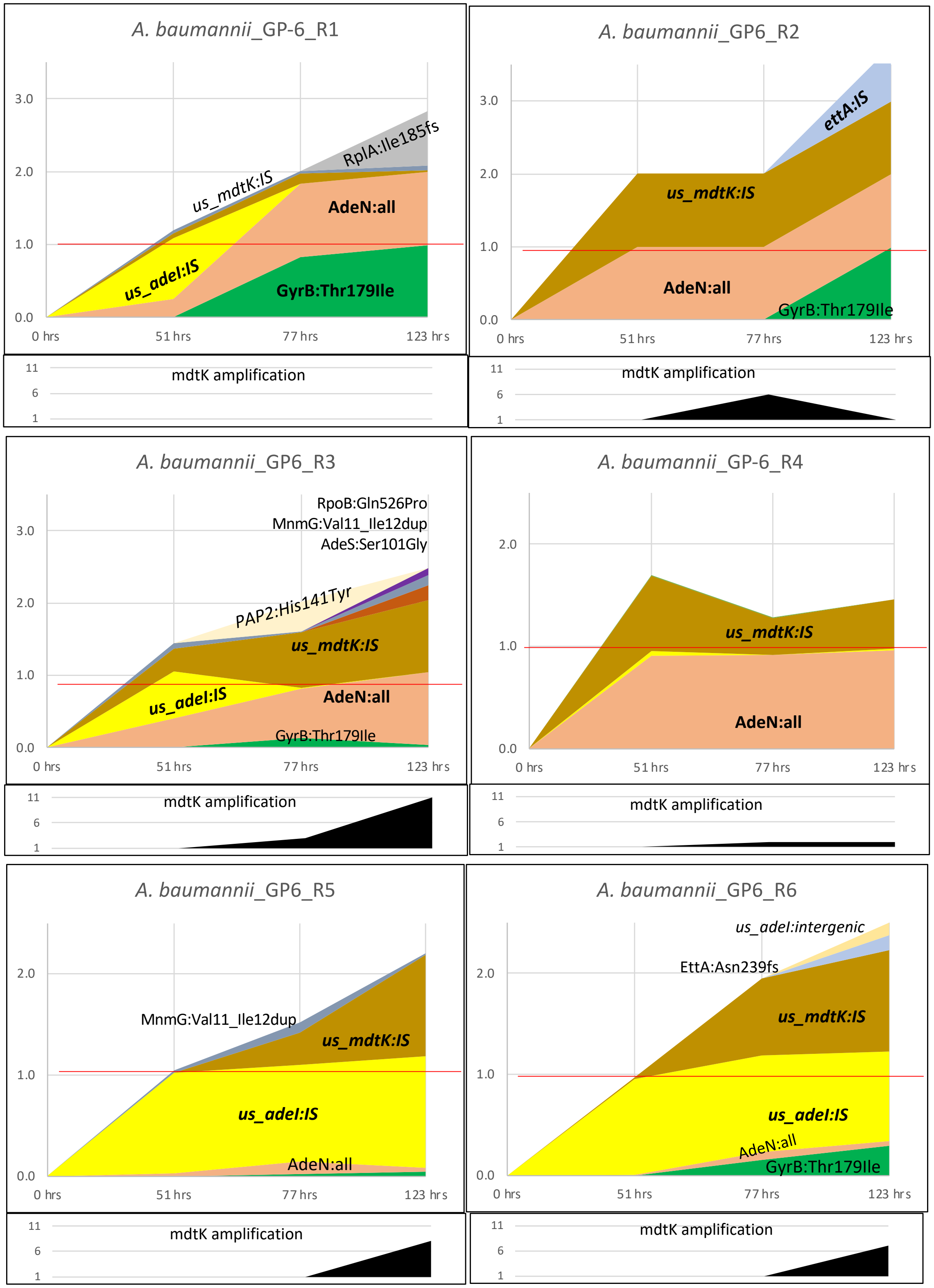

Cumulative area plots are shown for the most prominent mutational variants reaching relative abundance Amax ≥10% in at least one sample. Plots are shown for each of the six reactors R1-R6. The variant abundance data (from Supplementary Table S1 A and B) corresponding to each of the 6

reactors are plotted as the average number of corresponding mutations per cell (Y-axis) in the population vs time (X-axis). The red line (at average 1 mutation/genome) helps to emphasize the emergence of double and triple mutants. For each reactor, the upper plot shows relative abundances of SNVs and IS insertions. The lower plot shows the observed extent of amplification of MdtK locus, for which the relative abundance cannot be assessed in population WGS data.

*Gene names and corresponding functions are explained in the legend to Table 1 (and in Supplementary Table S1A with more details and tabular data).* A comparative analysis of these plots, which was only briefly described in the respective sub-sections of Results and Discussion, yields the following additional observations:

***Drug target modification***

*In E. coli (Figure S3A):*

**GyrB:** Among the 6 variants spanning 5 amino acid residues in *E. coli* GyrB, each of the two most prominent variants, GyrB:Thr165Ile and GyrB:Gly77Ser, dominated in one of the reactors reaching $A_{max}$=98-99%. The other three variants, GyrB:Thr165Asn, GyrB:Val167Glu and GyrB:Ala47Ser were less frequent reaching $A_{max}$ in the range of 4-14%. The only variant localized remotely from the ATP-binding site, GyrB:Cys476Phe, showed the lowest $A_{max}$=2%, and its relevance for GP-6 resistance remains questionable.

*In A. baumannii (Figure S3B):*

**GyrB:** Of the two detected variants of *A. baumannii* GyrB, the most prominent, GyrB:Thr179Ile (a full equivalent of *E.coli* GyrB:Thr165Ile), emerged in 5 out of 6 bioreactors reaching 99% abundance in two of them. The second minor variant, GyrB: Ala61Ser (matching *E.coli* GyrB:Ala47Ser), appeared only in 1 reactor reaching $A_{max}$=3% prior to being competed out by the GyrB:Thr179Ile variant.

***Efflux deregulation***

*In E. coli (Figure S3A):*

**AcrR:** In reactors R1 and R6, the aggregated abundance of AcrR:IS variants peaked at 54 and 75%, respectively, at an earlier stage of evolution while decreasing (down to 14% and 1%) by the end of the run. In one of them (R6), the AcrR:Thr5Asn variant emerged at the later stage competing out an earlier dominant AcrR:IS variant. Overall, missense mutations in the same position (AcrR:Thr5Asn and AcrR:Thr5Ile) emerged independently in three bioreactors (R1, R5, R6), reaching $A_{max}$~36-99%. Quite likely, missense mutations at Thr5 position (in the DNA binding domain of AcrR) also lead to at least partial loss of AcrR function as a repressor of *acrR<>acrAB* divergon. Finally, four distinct SNVs were detected in the intergenic *acrR<>acrAB* d region of the overlapping in three reactors (R1, R3, R5) reaching $A_{max}$~100% in one of them (R3).

**MarR:** Several intergenic mutational events were observed upstream of *marR* gene likely leading to derepression of *marRA* operon and, ultimately, to overexpression of AcrAB and TolC components of the *E. coli* efflux machinery. By extension, we consider a missense mutation MarR:Glu10Asp reaching $A_{max}$=28% in one of the reactors (R4) as potentially having a similar effect.

**MdtK:** We observed amplification of ~ 9 kB gene locus contains 2 genes upstream (*cfa* and *ribC*) and 6 genes downstream (ydhQ-W, mostly with unknown functions) of *mdtK* gene encoding a MATE family efflux transporter. Additional evidence implicating MdtK as a transporter contributing to efflux of GP-6 includes: (i) a detected intergenic mutation in *mdtK* upstream region potentially leading to upregulation of its expression; and (ii) similar genomic amplification of *A. baumannii* genomic locus harboring *mdtK* gene (see below) as the only common gene with the amplified locus in E. coli.

*In A. baumannii (Figure S3B):*

**AdeIJK:** Disruptive mutations (stop-gain, frameshift and IS insertions) in *adeN* gene encoding a repressor of a*deIJK* operon were observed in all 6 reactors rapidly expanding and sustaining at 100% aggregated abundance in 4 of them (R1-R4). Second, we observed two instances of ISAba1 insertions in the coding and intergenic regions of a gene (AUO97b_02717, a putative membrane-associated phospholipid phosphatase, PAP2-like) localized upstream of AdeIJK efflux pump encoding operon. They emerged at an early stage of evolution in 4 reactors (R1, R3, R5 and R6) reaching 65-100% abundance but sustaining in only two of them (R5 and R6) being competed out by AdeN:IS variants in the other two (R1 and R3). Of the two additional sporadic variants detected in the same locus (each at a single time point in a single reactor), a mutation in the intergenic region upstream of *adeIJK* operon may lead to its derepression (as in case of *adeN* disruptive mutations), while functional implications of a missense mutation His141Tyr in AUO97b_02717 (upstream of *adeIJK* operon) are unclear.

**AdeAB:** Two missense mutations in AdeS histidine kinase from AdeRS two-component system regulating AdeAB(C)[1] efflux pump were detected in R3. They emerged at the last stage of the evolution experiment reaching $A_{max}$=20% in case of the AdeS:Ser101Gly variant. Such variants were much more prominent in the previous CIP study (1), showing a broader repertoire of missense mutations (total of 7), more independent occurrences (total of 8) and higher $A_{max}$=65% (Table 1). Mutations in the *adeS* gene were previously observed as adaptation to antibiotic stress via AdeABC overexpression.

**MdtK:** We observed: (i) 3-11x amplification of ~11 Kb genomic locus containing *mdtK* gene in 5 out of 6 reactors*;* and (ii) insertion of *ISAba1* element in the gene immediately upstream of *mdtK* in all 6 reactors and reaching $A_{max}$~100% abundance in 5 of them; and (iii) deletion of ~2.5 kB genomic locus upstream of *mdtK* gene.

***Other potentially significant variants.***

<u>*In E. coli (Figure S3A)*</u>:

**Rob:** 3 mutational variants in the C-terminal domain of Rob transcriptional regulator: Rob:Ala201Thr, Rob:Gln194Pro and Rob:Cys147Tyr, the latter reaching 35% abundance.

**RpoC**: Gly336Ser variant ($A_{max}$=75% in R1) affecting β'-subunit of DNA-directed RNA polymerase complex (RNAP). While the impact of this mutation was not characterized, some mutational variants in RNAP β-subunit (RpoB) were implicated with CIP resistance and upregulation of *mdtK* gene (2). The latter effect, if phenocopied by RpoC: Gly336Ser variant could indeed impact GP6 resistance.

**Lon:** An insertion of IS186B mobile element in the upstream noncoding region of the *lon* gene was detected in reactors R1 and R2, in the latter reaching $A_{max}$=77%.

**YadG**: A functional significance of a frameshift variant YadG:Val83fs ($A_{max}$=96% in reactor R5) is unlikely since, being a singleton, it may be just a "hitchhiker" reaching high abundance due to accidental coupling with a genuine driver mutation. Such interpretation would be consistent with the observation that R5 is fully dominated by a single GyrB:Thr165Ile variant ($A_{max}$=100%). Although this variant outcompeted another early-stage double mutant, GyrB:Thr165Ile/AcrR:Thr5Ile, it is more likely due to the *marRAB locus duplication also identified in this clone (see Supplementary Table S2A) than to the mutation in yadG gene per se.*

<u>*In A. baumannii (Figure S3B):*</u>

**EttA**: disruption of ettA gene encoding energy-dependent translational throttle A

---

[1] Note that many strains of *A. baumannii*, including ATCC17978 contain only two genes encoding AdeAB but not AdeC components of this, otherwise functionally active, efflux pump. The nature of the alternative third component is under investigation (*Dr. A. Kumar, personal communication*).

emerged in two reactors (R2 and R6) as two distinct variants, EttA:Asn239fs; EttA:ISAba. The latter variant reached $A_{max}$=76% in the last stage of evolution on the background of a triple mutant GyrB:Thr179Ile/AdeN:ISAba1/*us_mdtK*:ISAba1.

**RplA:** of the 3 distinct frameshift variants in the gene *rplA* encoding ribosomal protein L1, one variant RplA:Ile185fs reached $A_{max}$=76% in the background of a double mutant GyrB:Thr179Ile/AdeN:ISAba1.

**MnmG:** a low frequency (Amax=15%) in-frame insertion in *mnmG* gene (involved in tRNA modification) leading to a Val11-Ile12 duplication was observed in 4 reactors.

**RpoB:** two low frequency variants ($A_{max}$=10%) were observed in RNAP β-subunit, RpoB:Gln526Pro and in-frame insertion/replacement variant RpoB:Asn777_Val778ins[ThrCysIleAsnGlnAsn]. A potential significance of these relatively minor variants is supported by the already mentioned precedents of RpoB mutations (2) and, indirectly, by the emergence of RpoC:Gly336Ser variant during evolution of GP6 resistance in *E.coli*.

**Supplementary Figure S4.**

**Genomic rearrangements leading to *mdtK* gene overexpression.**

**A:** Amplifications (amp) and deletions (del) in genomic loci harboring *mdtK* multidrug efflux transporter gene detected in evolving populations (left) and in 5 selected clones (right) of *E. coli* and *A. baumanni* analyzed by RT-qPCR.

**Genomic rearrangements in GP6 experimental evolution leading to MdtK overexpression**

| Evolving populations samples (WGS) | | | | | Clones (WGS+Nanopore): RT-qPCR(MdtK-OE); MIC(GP6) | | | |
|---|---|---|---|---|---|---|---|---|
| Sample | Copy (Ave) | From | To | Type | Clone | Variant | OE Log2FC | MIC Log2FC |
| **in *E. coli*** | | | | | | | | |
| **1F** | ~5x | 1,691,374 | 1,747,336 | Amp | | | | |
| **2F** | ~1x | - | - | WT | **2F4** | 1x(MdtK:us_del)+Acr:IS | 4.5 | 2 |
| **3F** | ~4.5x | 1,726,732 | 1,734,752 | Amp | | | | |
| **4F** | ~12x | 1,734,868 | 1,743,493 | Amp | | | | |
| **5F** | ~12x | 1,736,523 | 1,746,503 | Amp | | | | |
| **6F** | ~17x | 1,721,962 | 1,741,383 | Amp | **6F7** | 15xMdtK+AcrR:T5N | 2.8 | 4 |
| *(MdtK reference)* | | 1,737,714 | 1,739,087 | NA | | | | |
| **in *A. baumannii*** | | | | | | | | |
| **1F** | ~1x | - | - | WT | **1F1** | 1x(MdtK:us_IS)+AdeN:IS | 2.0 | 3 |
| **2D** | ~6x | 1,521,679 | 1,534,791 | Amp | | | | |
| **3D** | ~2.5x | 1,521,635 | 1,550,459 | Amp | **3D2** | 9x(MdtK:us_IS)+AdeN:IS | 5.9 | 5 |
| **4F** | ~0.05x | 1,518,115 | 1,521,507 | Del | **4F10** | 1x(MdtK:us_del)+AdeN:E198* | 5.5 | 5 |
| **5F** | ~7x | 1,521,635 | 1,535,875 | Amp | | | | |
| **6F** | ~5x | 1,521,331 | 1,535,153 | Amp | | | | |
| *(MdtK reference)* | | 1,521,873 | 1,523,168 | NA | | | | |

The extent of *mdtK* gene overexpression (OE) along with changes in $MIC^{GP6}$ are expressed as $\log_2$(FC)

**B:** Genomic rearrangements in the respective clones established by nanopore sequencing.

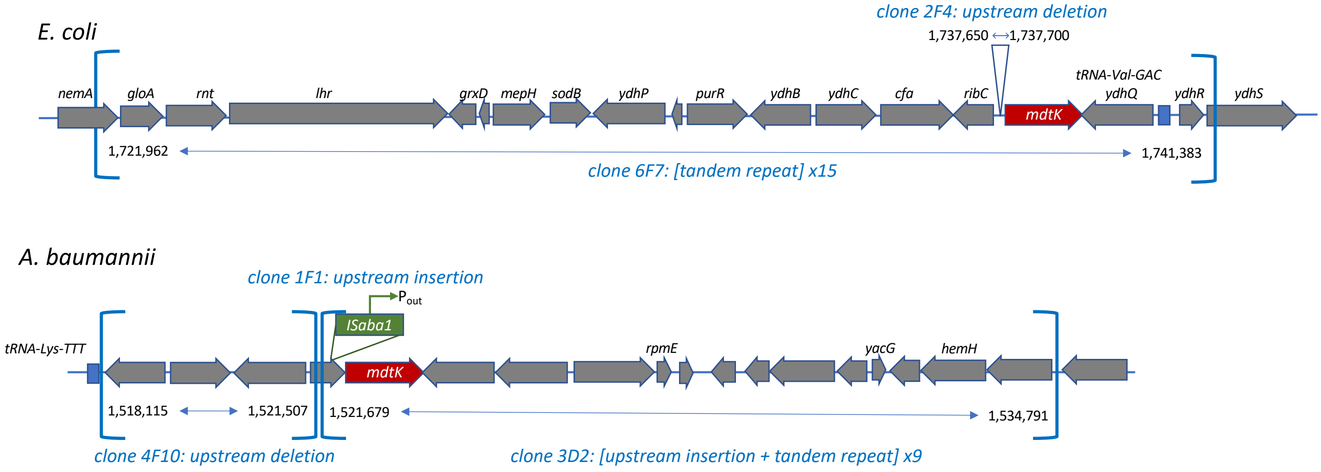

**Supplementary Figure S4 (continued).**

**C:** Testing of *mdtK* gene overexpression by RT-qPCR in selected *E. coli* clones harboring genomic rearrangements in *mdtK*-containing chromosomal locus: 50 bp upstream deletion (clone 2F4) or 15x locus amplification (clone 6F7). Clone 5C2 (carrying the same AcrR:IS5 variant as clone 2F4 but an intact *mdtK*-locus) and the unevolved parental strain (WT) are used as a negative control group.

*(Details of the experimental and statistical analyses are provided in Supplementary Metghods)*

**RT-qPCR analysis of mdtK expression in selected GP6-evolved *E. coli* clones vs WT**

| Sample | $C_t$ (gyrB) | $C_t$ (mdtK) | $\Delta C_t$ (mdtK-gyrB) | Average $\Delta C_t$ (mdtK-gyrB) | $\Delta\Delta C_t$ (relative to WT) | **$log_2$(FC) expression (realtive to WT** | **SD** | P-values one-way ANOVA with Tukey's HSD vs: | | |
|---|---|---|---|---|---|---|---|---|---|---|
| | | | | | | | | **2F4** | **5C2** | **6F7** |
| **2F4** | 18.26 | 16.68 | -1.58 | -1.463 | -4.54 | **4.54** | 0.15 | | | |
| | 17.75 | 16.45 | -1.30 | | | | | | | |
| | 17.93 | 16.42 | -1.51 | | | | | | | |
| **5C2** | 17.10 | 19.68 | 2.58 | 2.790 | -0.29 | **0.29** | 0.19 | 1.0E-07 | - | |
| | 17.16 | 20.02 | 2.86 | | | | | | | |
| | 17.42 | 20.35 | 2.93 | | | | | | | |
| **6F7** | 17.55 | 17.79 | 0.24 | 0.303 | -2.78 | **2.78** | 0.37 | 1.2E-04 | 9.6E-06 | - |
| | 18.04 | 18.01 | -0.03 | | | | | | | |
| | 17.56 | 18.26 | 0.70 | | | | | | | |
| **WT** | 17.55 | 20.37 | 2.82 | 3.080 | 0.00 | **0.00** | 0.25 | 1.0E-07 | 5.3E-01 | 4.2E-06 |
| | 17.74 | 21.06 | 3.32 | | | | | | | |
| | 17.75 | 20.85 | 3.10 | | | | | | | |

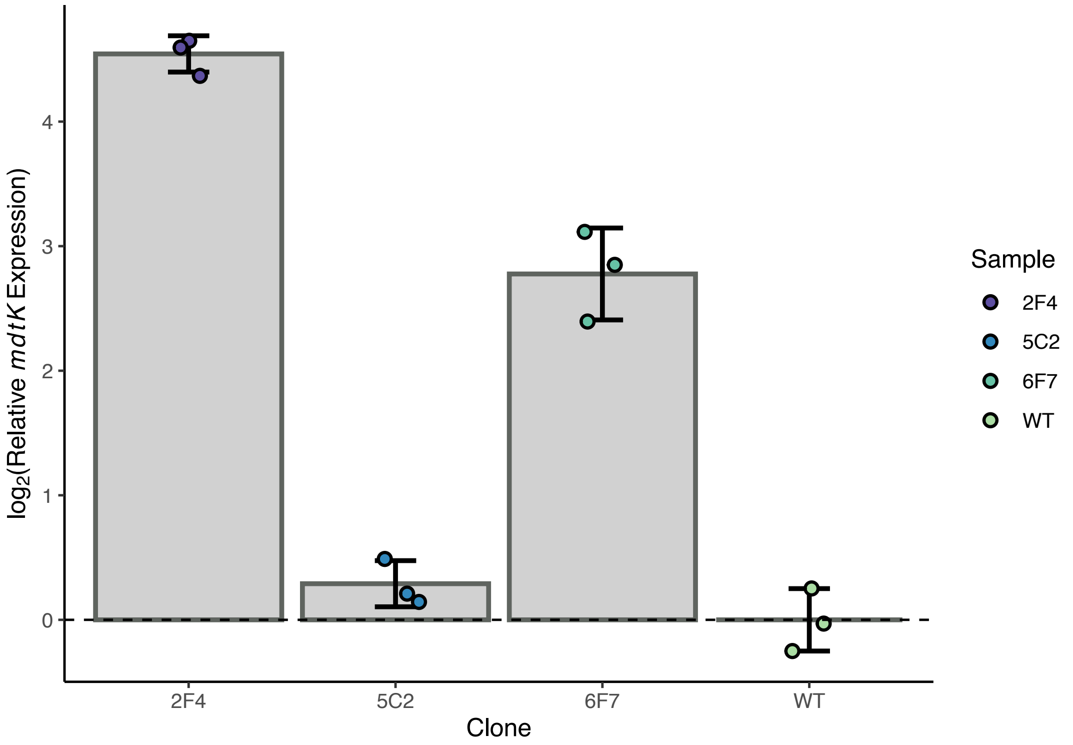

**Supplementary Figure S4 (continued).**

**D:** Testing of *mdtK* gene overexpression by RT-qPCR in selected *A. baumannii* clones harboring genomic rearrangements in *mdtK*-containing chromosomal locus: upstream IS-insertion (clone 1F1); upstream IS-insertion combined with 9x locus amplification (clone 2D2); or ~3.5 kB upstream deletion (clone 4F10). Clone 1F2 (carrying the same AdeN:IS variant as the clone 1F but an intact *mdtK*-locus) and the unevolved parental strain (WT) are used as a negative control group.
*(Details of the experimental and statistical analyses are provided in Supplementary Metghods)*

**RT-qPCR analysis of mdtK expression in selected GP6-evolved *A. baumannii* clones vs WT**

| Sample | $C_t$ (gyrB) | $C_t$ (mdtK) | $\Delta C_t$ (mdtK-gyrB) | Average $\Delta C_t$ (mdtK-gyrB) | $\Delta\Delta C_t$ (relative to WT) | **$log_2$(FC) expression (realtive to WT** | **SD** | P-values one-way ANOVA with Tukey's HSD vs: | | | |
|---|---|---|---|---|---|---|---|---|---|---|---|
| **1F1** | 19.1 | 19.58 | 0.48 | -0.253 | -1.96 | **1.96** | 0.64 | **1F1** | **1F2** | **3D2** | **4F10** |
| | 18.59 | 17.93 | -0.66 | | | | | | | | |
| | 18.49 | 17.91 | -0.58 | | | | | | | | |
| **1F2** | 18.25 | 20.24 | 1.99 | 2.140 | 0.44 | **-0.44** | 0.18 | 4.0E-03 | - | | |
| | 18.15 | 20.49 | 2.34 | | | | | | | | |
| | 18.33 | 20.42 | 2.09 | | | | | | | | |
| **3D2** | 18.44 | 13.71 | -4.73 | -4.195 | -5.90 | **5.90** | 0.76 | 2.4E-04 | 4.9E-06 | - | |
| | 18.42 | 14.76 | -3.66 | | | | | | | | |
| **4F10** | 18.4 | 14.35 | -4.05 | -3.810 | -5.51 | **5.51** | 0.67 | 2.2E-04 | 3.2E-06 | 9.4E-01 | - |
| | 18.33 | 15.28 | -3.05 | | | | | | | | |
| | 18.31 | 13.98 | -4.33 | | | | | | | | |
| **WT** | 18.55 | 20.56 | 2.01 | 1.703 | 0.00 | **0.00** | 0.52 | 1.4E-02 | 8.7E-01 | 8.9E-06 | 6.2E-06 |
| | 19.28 | 20.38 | 1.10 | | | | | | | | |
| | 18.13 | 20.13 | 2.00 | | | | | | | | |

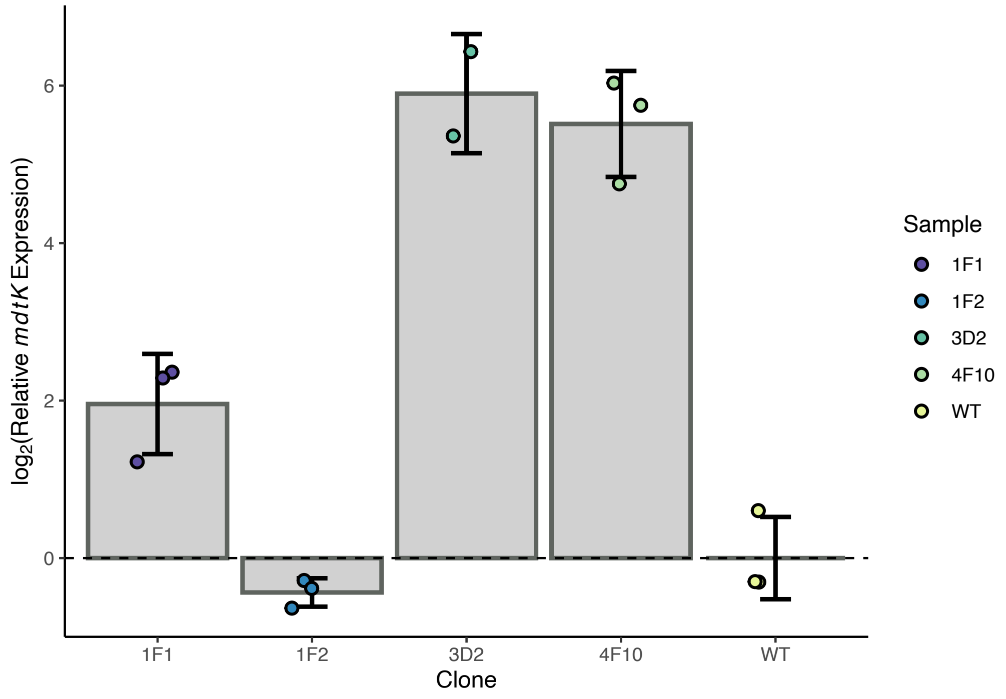

Supplementary figures

**Supplementary Figure S5.**

Structure of Rob transcriptional regulator complex with DNA (PDB:1D5Y (3)). Residues with variants observed in GP-6 morbidostat experiments are colored red.

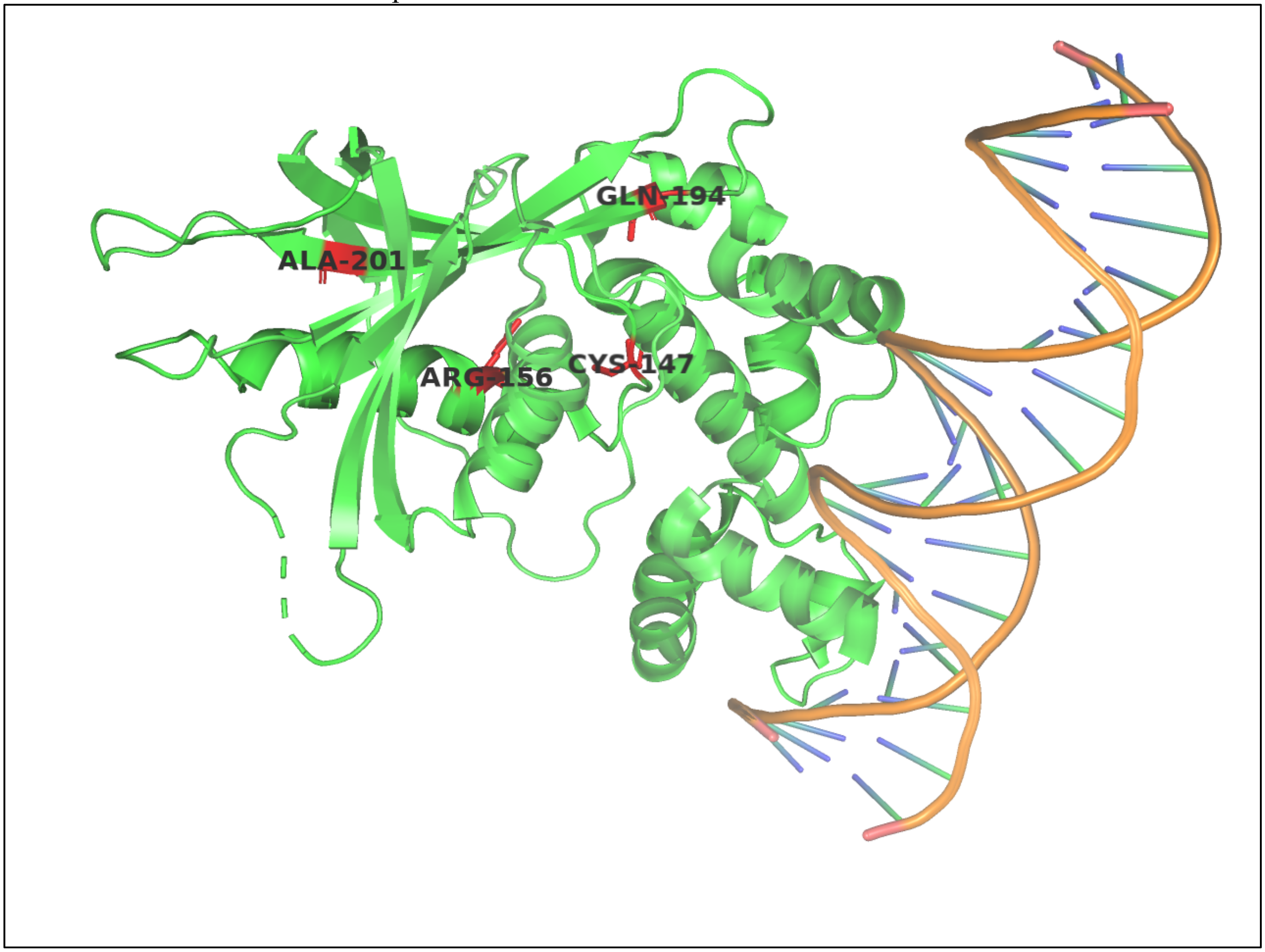
