## Supplemental methods for "Two Classes of DNA Gyrase Inhibitors Elicit Distinct Evolutionary Trajectories Toward Resistance in Gram-Negative Pathogens"

**Supplementary Methods**

**Supplementary Methods**

### **Morbidostat setup and programming**

Our implementation of the morbidostat approach was described in our previous studies with CIP and triclosan (1,2). Briefly, the morbidostat is a computer-controlled chemostat-like continuous culturing bioreactor where density of bacterial culture is controlled by varying an antibiotic concertation in media. Automated dilutions switch between addition of; (i) drug-containing media (increase of a drug concentration) when bacterial cultures are growing faster than dilution rate, and (ii) drug-free media (decrease of a drug concentration) when the growth rate slows down due to excessive drug pressure. This algorithm enables a gradually growing selective pressure driving the evolution of higher drug resistance. The detailed description of morbidostat implementation is provided on GitHub (<https://github.com/sleyn/morbidostat_construction>). The main characteristic features of our implementation include:

1) Six glass culture tubes containing 20 mL of media are used as bioreactors equipped with stirring bars for culture agitation and three needles for: (a) input of media and air, (b) waste removal and (c) sample collection.
2) Laser beam intensity is used to measure culture turbidity (optical density or OD at ~635 nm). The laser, photodiode and culture tube are installed in the 3D-printed rack.
3) A miniature air pump provides aeration and maintains air pressure for liquid displacement fitted with 0.22 µm air filters for sterility.
4) Drug-containing media and drug-free media are fed from two bottles connected via autoclavable silicon tubing to computer-controlled peristaltic pumps, each for one media type, followed by an array of 12 valves (2 valves per reactor, one for controlling flow of drug-free media and one – for drug-containing media).
5) The excess volume from the reactors (upon each dilution cycle) media is displaced to waste bottles by the constant air flow.
6) The temperature in morbidostat chamber is kept at 37^o^C with a heater controlled by thermal sensor.
7) All logic operations are controlled by an Arduino Mega 2560 Rev3 board. A user interface is implemented on PC using MegunoLink software (v.1.32.20005.0105; <https://www.megunolink.com/>).

The program logics and parameters of automated dilutions were as follows:

1) Regular dilutions with one or the other media (20% by volume) are made over programmable time intervals (cycle time, CT).
2) To select a dilution mode, OD by the end of each cycle is compared with two thresholds: lower threshold (LT) and drug threshold (DT).
3) At LT ≤OD <DT (as in the beginning of the run or after growth deceleration upon reaching drug concentrations above the level of acquired resistance), dilutions are made with drug-free media using CT=15 min (which could be manually adjusted up to 20-25 min upon overall fitness loss at higher drug concentrations)
4) At OD ≥ DT, the selection of the media for every dilution cycle (CT=15 min by default) is made based on the following logics:
   - if OD_1_ ≥OD_0_ where OD_1_ is OD at the end of the current cycle and OD_0_ is OD at the end of the previous cycle (e.g. when the culture growth rate is faster than the dilution rate), the dilution is made by the drug-containing media;
   - Otherwise, (if OD_1_ <OD_0,_ when the culture growth rate is slower than dilution rate), the dilution is made by the drug-free media.
5) At OD < LT (if it happens at poorly tolerable spike in drug concentration), the culture outgrowth was enabled via an automated increase of CT to 60 min with dilutions by the drug-free media (until reaching OD ≥LT).

### **Sequencing data analysis, variant calling and ranking**

The raw reads were processed as previously described (2). Briefly, upon adapter and quality trimming (BBDuk from BBTools suite v. 38.42, <https://sourceforge.net/projects/bbmap/>), the reads were aligned to reference genomes (BWA MEM v0.7.17 (3)). LoFreq Viterbi module was used to refine alignment near indel regions (4). Base Quality Score Recalibration was made by Genome Analysis Toolkit (GATK) modules BaseRecalibrator and ApplyBQSR (5). Sites with high frequency variants were masked from BaseRecalibrator by calling variants in the alignment down-sampled to ~ 50x coverage with Picard tools v.2.2.1 (<https://broadinstitute.github.io/picard/>) based on the coverage estimates by mosdepth v. 0.2.6 (6). The VCF file for BaseRecalibrator “--known_sites” option was produced by GATK HaplotypeCaller. All other SAM and BAM files manipulations (sorting, indexing, merging and splitting) were performed with Samtools v1.9 (3). SNP and indels were identified using LoFreq v2.1.3 (4).

Insertion sequence (IS) elements rearrangements were identified by a developed iJump tool (<https://github.com/sleyn/ijump>). IS elements in the reference genomes were predicted using BLAST against ISFinder database (7). Predicted effects of mutations were assigned with SnpEff v4.3 (8). VCF files manipulations were performed with bcftools v1.3 (9). Copy number variation (CNV) were predicted by CNOGPro package v1.1 for R (10). Reference genomes were downloaded from PATRIC database (11). Statistics of read alignments against reference genomes is in Supplementary Table S1C for population samples and is in Supplementary Table S2C for clone samples. Reference genome IDs are 679895.18 for *E. coli* BW25113 and 400667.82 for *A. baumannii* ATCC17978. WGS data and assembly of all six unevolved clones (A1 – A6) including variations compared to reference genomes were reported in the previous study (2). Nanopore reads were base-called using Guppy v.5.0.16 and aligned to reference with minimap2 v.2.17 (12). Large rearrangements were verified using Ribbon tool (13).

Ranking of the observed variants as significant was performed based on their statistics across all six reactors as described (2) and illustrated in Supplementary Tables S1A and B that provide complete lists of observed variants combing SNVs, short indels, IS inserts and CNV data. Their ranking is reflected in the column “Category” of both tables, which is used to apply filters limiting the display to highly ranked (potential drivers and strongly coupled mutational events). Our approach to ranking, after exclusion of technically unreliable calls (frequency <2%, imperfect repeats, etc.) as well as “preexisting” variants and synonymous mutations was gene-centered based on two major (complementary) features. First, we prioritize variants reaching high frequency (or maximal abundance, in a population sample weighted on the total site abundance, ***A_max_***) in at least one sample across all 6 reactors. Then, for all genes implicated by at least one variant with ***A_max_*** ≥10% (coding sequences and their upstream noncoding regions, and, in case of IS-inserts or large deletions, including upstream coding regions) we consider all variants (non-preexisting, non-synonymous and with ***A_max_***≥2%). The second feature for further ranking of these initially prioritized genes, reflects the overall number of independently occurring variants per gene (***N_all_***). The latter combines a total number of distinct variants (***V***) with the number of reactors where each variant independently arises (***R***). Of those, the first parameter (***V***) is typically more significant, especially for disruptive mutations in coding regions (such as frameshifts, stop-gain and IS inserts). Therefore, we additionally prioritize the genes with high values of V>3, even with 2%≤***A_max_***<10% for all distinct variants, as long as their total abundance exceeds 10% in at least one sample. This approach typically implicates up to 8-10 genes with ***A_max_***≥50% and *N_all_*≥3 (up to 20 for disruptive mutations) as illustrated in Table 1. A functional significance of up to 5-6 genes with lower rank (smaller values of one or both parameters) remains questionable, and, typically, not supported by clonal analysis. In contrast to SNVs and even IS-inserts (where abundance determination is intrinsically less accurate), CNVs (large deletions and amplifications) cannot be accurately estimated in population WGS data. Besides, they cannot be unambiguously assigned to a specific gene, as typically cover from 3 to more than 30 genes (see Supplementary Tables S1A and B). Therefore, the prioritization (and interpretation) of the observed CNV events relies on the comparison with a list of genes implicated by other types of events (as for *mdtK* and *marR* genes in this study). Identification of mutations in clones is straightforward. The observed variants typically match those from respective population data (although in rare cases additional mutations are observed), and their calculated abundance is usually >90% (see Supplementary Tables S2A and B).

Applying these criteria to both WGS datasets allowed us to define the list of significant variants (excluding CNVs): (i) for *E. coli*, 34 distinct events in 14 genes; and (ii) for *A. baumannii*, 30 distinct events in 10 genes (displayed in the **Supplementary Tables S1A and B** after applying respective filters).

### **Population WGS data deconvolution for evolutionary dynamics and clonal analysis**

The overall abundance of all (even highly ranked) mutations in a sample commonly exceeds 100%, especially at the later stages of experimental evolution pointing to the emergence of double and triple mutants (more characteristic of *A. baumannii* than *E. coli*, see Supplementary Figure 3). Although, the accurate assignment of these mutations to clonal subpopulations in many cases is nearly impossible, we used an approximate deconvolution of population samples for assessment of trends in evolutionary dynamics and optimization of clone isolation strategy aimed to maximize the coverage of the major mutational variants and combinations thereof.

We used Mixed Integer Programming to deconvolute population sequencing data represented by a matrix of variant frequencies (***F***) where rows are variants, columns are sampling time points and values represent variant frequencies in the population raging from 0 to 100% (as in *Supplementary Tables S1 A and B*). Since population samples are composed of one or more competing subpopulations that emerge and expand or wash out, each variant frequency (***f***) is a sum of frequencies of subpopulations where this variant is present:

$$f_{n}^{k}= \sum_{i=1}^{m} s_{i}^{k}p_{ni}^{k}$$

where ***f_n_^k^*** is a frequency of an *n-*th observed variant in a *k-*th sample; *s_i_^k^ –* frequency of an *i*-th subpopulation in the *k-*th sample, *p_ni_^k^* – a dummy variable, which is equal to 1 if the *n*-th variant *n* is present or 0 if it is absent in the *i*-th subpopulation in the *k-*th sample. Then the matrix of variant frequencies ***F*** can be computed as matrix multiplication:

$$P \cdot S=F$$

where *P* is a binary matrix of dummy variables with rows representing variants and columns representing subpopulations; and *S –* a matrix of subpopulation frequencies with rows representing subpopulations and columns representing population samples. The deconvolution of populations is formalized as an optimization problem to find values of matrices *P* and *S* given the following constrains:

1) Values of matrix *P* are binary.
2) Values of matrix *S* have boundaries 0 and 100%.
3) Sum of each column of matrix *S* should be within the range from 0 to 100%.

The algorithm was implemented using PuLP toolkit of Python 3 (14) and Gurobi solver (15). The details of the algorithm and its implementation could be found at the GitHub page (<https://github.com/sleyn/subpop_decompose_mip>). The obtained subpopulation frequencies were used to choose the samples with the largest representation of the most prominent subpopulations (with s≥20%) that would allow us to maximize the odds and minimize the redundancy in their representation in clonal analysis.

### **RNA isolation and RT-qPCR**

RNA isolation from flash-frozen pellets was performed as described using a modified protocol of (16) with minor modifications. Briefly, cell pellets were resuspended in lysis buffer ( NaCl, 140 mM; EDTA, 14 mM; SDS, 6 % w/v) and mixed with Phenol_Chloroform: Iodoacetic acid (125: 24: 1, pH 4.5). Glass beads (acid washed, 500 uL) were added to each sample before being homogenized using a Bead Ruptor 12 (Omni Inc) for 2 minutes at 6 m/s 2 times. Subsequently, the samples were centrifuged (15 minutes, 16000 *g*, 4 °C) and the aqueous phase collected. Crude RNA was precipitated from this phase using sodium acetate (0.3 M final concentration) and isopropanol overnight at -20 °C. The precipitates from overnight were collected by centrifugation (30 minutes, 16000 *g*, 4 °C), washed 2 times in ice cold ethanol (70 %), dried in a laminar flow hood, and resuspended in nuclease free water. Coprecipitated DNA was removed using Baseline-ZERO^TM^ DNase (Lucigen) following the manufacturers protocol. Samples were further purified using the MEGAclear^TM^ Transcription Clean-Up Kit (Invitrogen). RNA concentration and quality was assessed via both nanodrop, using A260/280 and A260/230 ratios, and 1 % TAE agarose gel to monitor ribosomal RNA integrity. For samples which exhibited poor A260/A230 ratios, an additional round of ethanol precipitation was performed using the GlycoBlue^TM^ coprecipitant kit (Invitrogen), following the manufacturers instructions. Isolated RNA was frozen in liquid nitrogen and stored at -80 °C.

Complementary DNA was prepared from the isolated RNA using the High Capacity cDNA Reverse Transcription Kit (Thermo Fisher Scientific) following the manufacturers direction, with 200 ng of cDNA being prepared per clone per replicate. qPCR reactions were prepared using the PowerTrack^TM^ SYBR^TM^ Green Master Mix (Thermo Fisher Scientific) with 10 ng of prepared cDNA being used per gene for each sample and a final primer concentration of 400 nM. The reactions were performed on a Bio-Rad CFX384 Touch Real-Time PCR System with the following conditions: 95 °C for 10 minutes; 40 cycles of 95 °C for 15 s, 55 °C for 60 s. Threshold cycle (Ct) values were assigned for both the target gene, *mdtK*, and the reference gene, *gyrB* within CFX Manager^TM^ (Bio-Rad). Relative expression of *mdtK* and the associated standard deviations calculated for each clone using the 2^-ΔΔCT^ method (17). One-way ANOVA with Tukey’s HSD post-hoc testing was performed in R. The primers used are shown below.

For *A. baumannii*

Target for the assessment of overexpression

*mdtK* FW primer:	ATTCCAGTGATTGTCCGCCA

*mdtk* RV primer:	ATGCTTCGGAATAGCCTCGG

Housekeeping control for normalization

*gyrB* FW primer:	GATGATGCGCGTGAAGGTTT

*gyrB* RV primer:	CATTGCTTGCTCTACCGCTG

For *E. coli*

Target for the assessment of overexpression

mdtK FW CTCTTTGGTCACGGACTGCT

mdtk RV TACGGGCAACCTGGAAGAAC

Housekeeping control for normalization

gyrB FW GAACAAAACGCCGATCCACC

gyrB RV GCACTTTCACGGAAACGACC
